# An integrated metagenome catalog reveals novel insights into the murine gut microbiome

**DOI:** 10.1101/528737

**Authors:** Till Robin Lesker, Abilash Chakravarthy, Eric. J.C. Gálvez, Ilias Lagkouvardos, John F. Baines, Thomas Clavel, Alexander Sczyrba, Alice C. McHardy, Till Strowig

**Author notes:** Corresponding author: Till Strowig.

## Abstract

The vast complexity of host-associated microbial ecosystems requires generation of host-specific gene catalogs to survey the functions and diversity of these communities. We generated a comprehensive resource, the integrated mouse gut metagenome catalog (iMGMC), comprising 4.6 million unique genes and 660 high-quality metagenome-assembled genomes (MAGs) linked to reconstructed full-length 16S rRNA gene sequences. iMGMC enables unprecedented coverage and taxonomic resolution, i.e. more than 89% of the identified taxa are not represented in any other databases. The tool (github.com/tillrobin/iMGMC) allowed characterizing the diversity and functions of prevalent and previously unknown microbial community members along the gastrointestinal tract. Moreover, we show that integration of MAGs and 16S rRNA gene data allows a more accurate prediction of functional profiles of communities than based on 16S rRNA amplicons alone. Integrated gene catalogs such as iMGMC are needed to enhance the resolution of numerous existing and future sequencing-based studies.

## Introduction

The gut microbiota is a dynamic and highly diverse microbial ecosystem that impacts the hosts physiology^1^. Culture-independent methods such as high-throughput sequencing have revolutionized experimental approaches to characterize and investigate these communities. Gene catalogs facilitate taxonomic and functional annotation of sequencing data, thereby maximizing insights gained from short-reads^2–5^. Moreover, they can provide higher resolution than less specific resources such as GenBank by including valuable metadata such as environment-specific variables. Typically, generation of reference gene catalogs involves sample-specific assembly, prediction of genes and dataset-wide clustering of gene entries to reduce redundancy. However, this approach results in reduced taxonomic resolution of gene entries, first due to clustering of highly related but distinct genes and second due to the lack of high-resolution taxonomic information for gene entries, which can be best obtained from marker genes, such as 16S rRNA genes for which large reference collections exist. Here we present a novel approach and corresponding computational workflow to construct integrated gene catalogs, resulting in a significant improvement of the taxonomic resolution of gene entries and providing valuable additional information such as linking genes to metagenome-assembled genomes (MAGs) and reconstructed full-length 16S rRNA genes. We applied this approach to construct an integrated mouse gut metagenome catalog (iMGMC) combining existing and newly sequenced metagenomic data. We chose this ecosystem as the mouse serves as foremost experimental model system to study microbiota-modulated human diseases, but the use of currently existing human gut gene catalogs is precluded due to the substantial differences in bacterial species and genes present in mice^6^.

## Results

### Construction of the integrated mouse gut metagenome catalog (iMGMC)

Pioneering work by others resulted in the construction of several gene catalogs, including a microbiome gene catalog from the mouse gut (hereon referred to as MGCv1) comprising 2.6 million non-redundant genes^4^. We developed a bioinformatic workflow that combines a global assembly strategy with binning of contigs to putative MAGs and with innovative linking of reconstructed 16S rRNA gene sequences to these MAGs (Figure 1A). This “All-in-One” assembly approach together with the subsequent binning enables maintaining complex information such as distribution of distinct contigs and bins over a large number of samples. We applied this approach to a previously published set of sequencing data included in MGCv1 (n = 190 mouse fecal samples) and increased the biological diversity by incorporating novel metagenomic data for 108 additional intestinal samples from a large number of commercial mouse providers and wild mice, including different gastrointestinal locations (see Table S1). This selection was based on the previous notion that the source of experimental mice and anatomic niches contribute to the variability between murine microbiome to a higher extent than other factors such as diet, genotype, housing laboratories or gender^4^. As a first step in the construction of iMG2C, 1.3 Tbp from 298 metagenomic sequencing libraries were assembled using Megahit^7^ in an “All-in-One” approach, resulting in 1.2 million contigs of length greater than 1000bp, with a total assembly size of 4.5 Gbp. Next, genes were identified with MetaGeneMark^8^, resulting in 4.6 million open reading frames (ORFs) of length greater than 100 bp, compared to 2.6 million ORFs in the MGCv1 (+77%) (Figure 1B). We tested the redundancy of these ORFs by clustering them with CD-Hit (95% identity at 90% coverage)^9^, which resulted in a reduction of only 2% of ORFs (n = 99,670) (data not shown). We considered this negligible compared to the 89% reduction in MGCv1 ^4^. Subsequently, contigs were binned using MetaBat^10^, resulting in 1,462 bins greater than 200 kbp (containing 87% of iMGMC entries). Subsequently, we defined 660 bins encoding 40% of all iMGMC entries as MAGs, based on the presence of established sets of bacterial marker genes and a quality threshold ≥80% (Figure 1C) ^11^. Notably, MGCv1 did not provide MAGs, as sample-specific assemblies were used, but rather less specific information referred to as “co-abundance groups” (CAGs), containing at least 700 genes. Comparison of the numbers of CAGs and genes in CAGs between iMGMC and MGCv1 revealed large increases in our resource (1,217 vs. 541 CAGs, 81% vs. 40% of genes, respectively) (Figure 1B).

**Figure 1:**
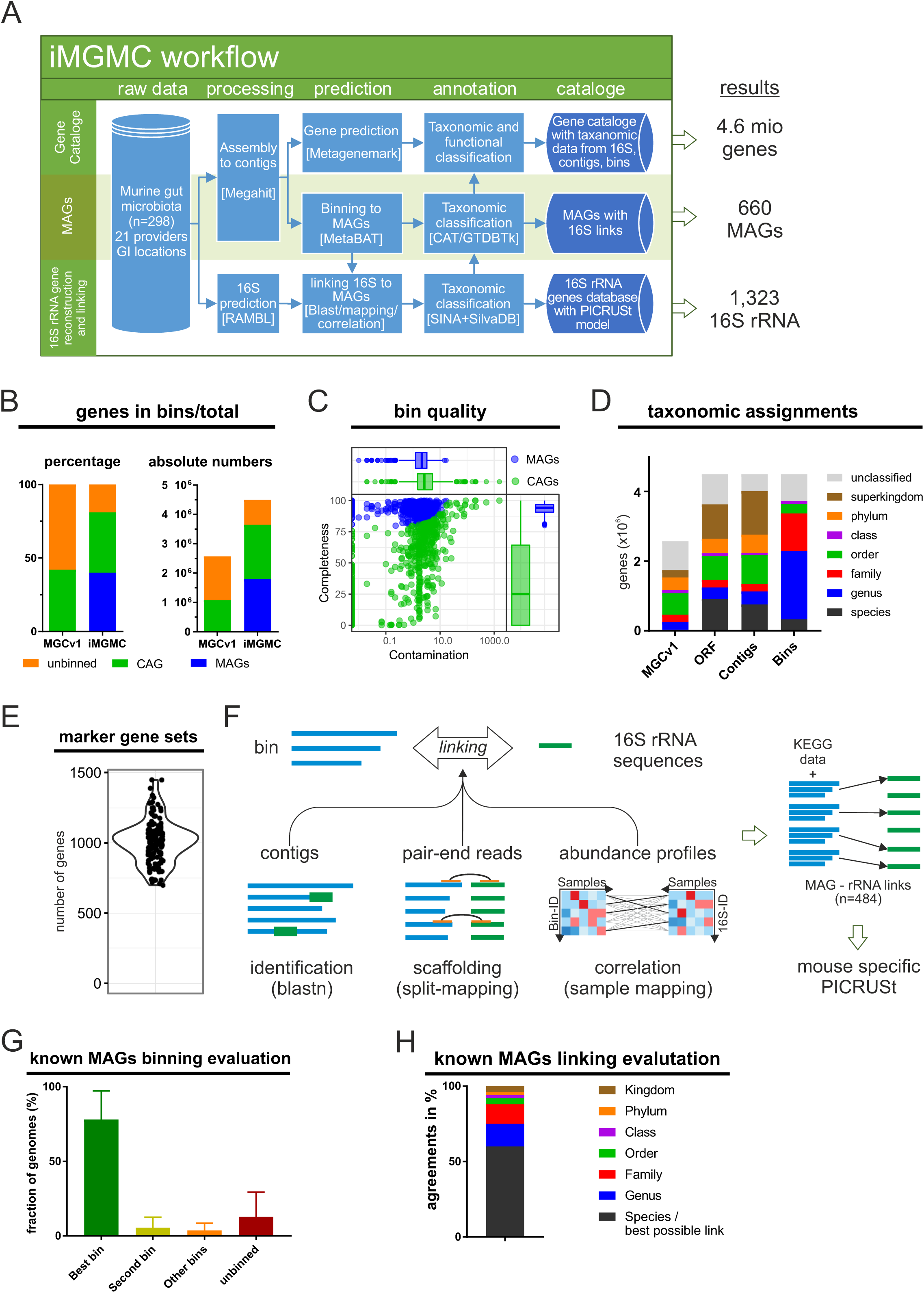
Generation and evaluation of the integrated mouse gut metagenome catalog (iMGMC) (A) Flowchart displaying the steps and bioinformatics tools (names in brackets) utilized for the generation of the iMGMC. This resource includes genes, metagenome assembled genomes (MAGs), 16S rRNA gene sequences and MAG-16S rRNA gene links. (B) Comparison of relative and total numbers of gene entries and their association to bins of different completeness between a previous mouse gut gene catalog (MGCv1)^4^ and iMGMC. Bins were defined as: i) co-abundance genomes (CAG) if they were larger than >= 200kbp lengths and contained ≥700 ORFs or: ii) MAGs if their quality (marker gene completeness – contamination) as determined by CheckM was ≥ 80%. (C) Quality determination of individual binned contigs by CheckM by analyzing marker gene completeness and contamination. Box plots display marker gene completeness and contamination of 660 MAGs and 802 CAGs, respectively. (D) Absolute numbers of gene entries colored according to the lowest possible taxonomic annotation of the ORF, contig or bin. Different taxonomic profilers were employed for classification: ORF: DIAMOND-BlastPp contigs: CAT (Contig annotation tool)p bins: GTDBTk (E) Number of genomes in dataset estimated using a marker gene set containing 139 genes. Each dot represents the copy number of the respective marker gene. (F) Overview of the methodology to link MAGs to 16S rRNA gene sequences by combining mapping-based and statistical approaches. Resulting linked pairs of MAGs and reconstructed 16S rRNA gene sequences were used together with KEGG annotations for construction of mouse gut specific PICRUSt predictions. (G) Evaluation of binning by calculating the fraction of recovered RefSeq genomes (threshold ≥ 50 % of genome present in contigs, n=57) in bins. (H) Evaluation of MAG / 16S rRNA gene linking by determining the taxonomic match between predicted and reference 16S rRNA gene sequence for those recovered RefSeq genomes with a MAG / 16S rRNA gene pair (n=47). See Figures S1, S2 and S3 for more details.

In addition to reconstructing bins including MAGs, we also assembled 16S rRNA genes, using the following approach that overcomes the limitation that 16S rRNA genes are typically not efficiently recovered in standard assemblies, due to their highly conserved regions^12^: Using RAMBL^13^, we reconstructed 1,323 full-length, unique 16S rRNA gene sequences, a number similar to the number of genomes (n=1,068) predicted based on the presence of 139 distinct marker genes in the iMGMC assembly using Anvi’o (Figure 1E)^14^. We postulated that linking 16S rRNA genes to bins and MAGs after assembly would allow efficient integration of these complementary pieces of information, thereby improving the taxonomic assignment of MAGs. However, no high-throughput method currently exists for creating such links. Hence, we developed an integrated score combining mapping- and correlation-based associations to assign a 16S rRNA gene sequence to each bin and MAG (Figure 1F and S1). Briefly, we first identified all contigs containing reconstructed 16S rRNA gene sequences via BlastN^15^. Then, we searched for paired-end reads in which one read mapped to a reconstructed 16S rRNA gene sequence and the other to a contig. Finally, we remapped all libraries to the 1,462 bins and the 1,323 16S rRNA gene sequences to determine their relative abundances across all samples and used this data to estimate correlations between bins and 16S rRNA gene sequences using an abundance co-variance strategy^16^. This individual information was finally integrated using a novel approach (see Methods for details) to assign the reconstructed 16S rRNA genes to bins.

### Evaluation of iMGMC generation

The different steps underlying the construction of iMGMC were evaluated for their efficiency using those MAGs that had a highly related reference genome. These were specifically identified by mapping synthetic reads generated with BBMap from all 9,748 bacterial genomes available in the NCBI Assembly database (Version January 2017) against all bins and also the contigs that we were not able to bin (unbinned contigs) in iMGMC (see Methods for details). After read mapping, we evaluated the distribution of these genomes in our assembly and identified 57 genomes, which were recovered at least by 50% within binned and unbinned contigs. For these genomes, we recovered on average 79 ± 11% (mean ± s.d.) in our assembly, from which 78 ± 19% were found in the respective best/largest bin, while only 13 ± 17% were found in unbinned contigs (Figure 1G and S2). Thus, we considered our “All-in-one” assembly as good as other assembly strategies employed for large-scale MAG reconstruction^17^. The number of MAGs (n=660) would even be higher when using a quality threshold from an already published study (n=818, quality(CheckM): Completeness – 5x contamination ≥ 50%)^17^. We also evaluated the utility of the “All-in-one” assembly approach for another large dataset by processing metagenomic sequencing data from the pig microbiome. From 287 fecal samples (1,758 Gb) used to construct a previous reference gene catalog^5^, we obtained 12.2 Mio ORFs and 1,050 MAGs, representing a 58%- and 45 %-increase, respectively, compared with the original work (data not shown).

The MAG/16S rRNA gene pairs were evaluated using MAGs with linked 16S rRNA gene sequences for which reference genomes exist. Specifically, we identified genomes found in our assembly and the respective bins, followed by comparison of the known 16S rRNA gene sequences to the correspondingly predicted 16S rRNA gene sequences (Figure S3) (see Methods for detail). From the 47 identified genomes and respective bins, 28 agreed perfectly (100% sequence identity) between known and linked 16S rRNA gene, with an additional 7 matching taxonomic assignment down to the genus level. The remaining 12 genomes and bins disagreed at varying taxonomic levels (Figure 1H and S3). Statistical assessment of these results supported that our approach i) did not require 16S rRNA gene sequences within a MAG to successfully perform a matching linking and ii) performed better than a random assignment (P=0.074, Pearson’s Chi-squared test with Yates’ continuity correction). Hence, the proposed novel scoring scheme is with high confidence able to link MAGs and bins to corresponding reconstructed 16S rRNA genes, improving taxonomic resolution, though not in an error-free manner.

Thus, we created a novel type of resource which i) includes a gene catalogs that outperform previous versions and ii) includes novel information, i.e. MAGs, and 16S rRNA gene sequences, which are linked with each other.

### iMGMC reveals high prevalence of novel taxa in the mouse gut microbiota

Both metagenomic and cultivation-based studies showed that the gut microbiome of mice compared to human is composed of distinct bacterial species, of which many are yet uncultured and lack genomic information^4^,^6^. Analysis of our 660 reconstructed MAGs corroborates this notion, revealing that only 72 of them have closely related NCBI assemblies including other MAGs available (ANI > 95%) (Data in Table S1)^18^. A similar observation (only 137 known of 1,050 MAGs in total) was made for MAGs derived from the pig microbiome.

To construct a comprehensive phylogenetic tree of the mouse gut microbiota, we assigned MAGs (n=660) and closely related, previously sequenced genomes (n=64) into clusters (Figure 2). In line with previous reports^6^,^19^, our data analysis corroborates that the murine gut microbiome is overall dominated by the two main phyla *Firmicutes* (77% of MAGs / 73% of 16S rRNA gene sequences) and *Bacteroidetes* (14% / 18%)(Figure 2 and S4). Notably, *Bacteroidetes* included the second largest MAG cluster, namely the *Bacteroidales* S24-7 group (64% / 49%), recognized as being very abundant in the mouse gut, but for which only three reference genomes are available 6(new Microbiome paper). Strikingly, ≥13 % of MAGs were from phylogenetic groups (up to level of family) that completely lacked reference genomes in public databases (NCBI genomes RefSeq, not other MAGs), such as MAGs assigned to the *Clostridiales*-vadinBB660 group (n= 70) and *Mollicutes* RF9 (n=14) (Figure 2). Unsupervised clustering of MAG according to their functional potential (Figure S5) demonstrated that distinct taxonomic clusters such as *Clostridiales*-vadinBB660 group or the *Bacteroidales* S24-7 group represent functionally distinct microbes within the mouse microbiome (Figure S7) (new Microbiome paper).

**Figure 2:**
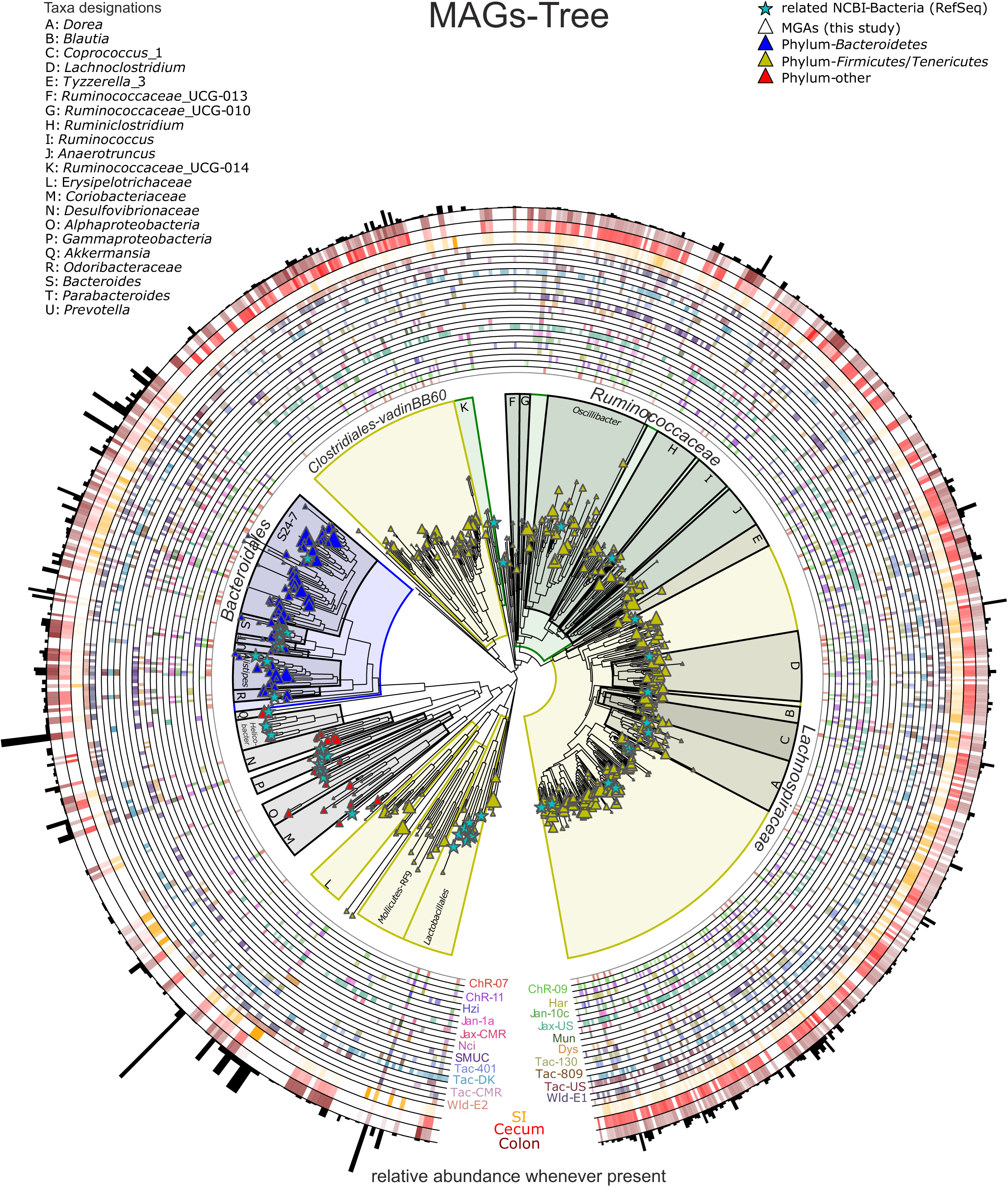
Phylogenetic tree of the 660 MAGs included in the iMGMC. MAGs are shown as triangles and 64 closely related, previously sequenced bacteria used for comparison as stars (genomes from NCBI refSeq with mapping rate >50% coverage). The color of triangles indicates their taxonomic association to different phyla and the size of triangles indicates the mean relative abundance in all iMGMC samples. The tree includes manually curated taxonomic assignments for most MAGs and the names of the taxonomic clusters are displayed in full or abbreviated in the tree. The inner rings show the relative abundance of the 660 MAGs in the 21 investigated mouse providers (threshold: 0.1%). The last three rings visualize the relative abundance of 469 of 660 MAGs at different anatomical sites (threshold: 0.1%, SI: small intestine). The outer bar plots show their respective maximal relative abundance.

Many additional undescribed bacteria were also identified after comparing the reconstructed 16S rRNA gene sequences to members of “16S ribosomal RNA (Bacteria and Archaea)” at the NCBI-database, with only 164 of 1,323 (12%) having at least a 97% identical match. A large fraction of these sequences were neither found in the SILVA SSU Ref v. 128 database (99% ident: 72% new, 97% ident: 45% new) nor in a recent 16S rRNA database established by target-specific environment sequencing^20^ (99% ident: 98% new, 97% ident: 93% new). Notably, while the MAGs represent a large fraction of the phylogenetic tree of the bacteria present in the mouse gut, several taxonomic groups were represented by 16S rRNA gene sequences, but underrepresented by MAGs, such as the family of *Prevotellaceae* (49 16S rRNA gene sequences / 3 MAGs), the class of *Bacilli* (81/10) as well as the phyla of *Proteobacteria* (67/24) and *Actinobacteria* (78/22) (Figure S4). Thus, our analysis identified taxonomic groups that are interesting novel targets for cultivation-dependent and -independent studies to extend our understanding of microbiome-modulated phenotypes in mouse models.

### Improved functional prediction via MAG/16S rRNA gene links in iMGMC

The establishment of databases of microbial reference genomes has spurred the development of computational approaches to simulate the functional profiles of metagenomes based on marker gene datasets such as 16S rRNA amplicon profiles^21^,^22^. However, the power of these approaches depends on the availability of sequenced microbial genomes from the respective environments to perform satisfactorily. Because of the existence of numerous bacterial species within murine gut communities that lack reference genomes, we hypothesized that the default PICRUSt-based predictions of mouse-associated metagenome functions are limited^21^. Thus, we constructed a mouse-optimized PICRUST version, employing the original PICRUSt algorithm in conjunction with the iMGMC data. Specifically, we used the MAGs with unique linked 16S rRNA sequences (n=484), as well as the 1,322 16S rRNA sequences from the iMGMC to create an extended genome resource for PICRUSt (PICRUSt-iMGMC) (Figure 3A, see methods for details). Comparison of Kegg Ortholog (KO) functional profiles predicted by the default and extended PICRUSt approach using 16S rRNA amplicon data from different gastrointestinal sites (n=50) for the corresponding shotgun metagenomic libraries (WGS) demonstrated a higher correlation to the WGS-based KO profiles for PICRUSt-iMGMC than PICRUSt-default predicted profiles (Pearson: 0.84 vs 0.68, +23%, Spearman: 0.84 vs 0.70, 21%) (Figure 3B and C). The highest correlations were observed for samples from the colon (Pearson: 0.86 vs. 0.67, Spearman: 0.87 vs. 0.72) (Figure S6). Similar improvements were obtained with distinct datasets not used for the construction of the catalog (Figure S7). The improved correlation of PICRUST-iMGMC largely derived from increased sensitivity, i.e. “true positive rates”, rather than decreased “false positive rates”, enabling the prediction of functionalities otherwise lost (Figure 3D and E). Even when mapping WGS data to the KEGG database with DIAMOND^23^ instead of to the iMGMC for generation of the KO reference profile, PICRUSt-iMGMC performed better than PICRUSt-default in predicting functional profiles (Figure S6 and S7). Finally, we evaluated whether combining the information of iMGMC with the genomes available in the KEGG database improved prediction. Strikingly, PICRUSt-iMGMC/KEGG did not perform better and the correlation with WGS data even decreased, suggesting that inclusion of related but divergent genomes reduces prediction accuracy (Figure S6 and S7). Hence, our resource enabled the development of ecosystem-specific PICRUSt models, i.e. optimized for the murine intestinal microbiome, with substantial improvement in the prediction of metagenomic functional profiles.

**Figure 3:**
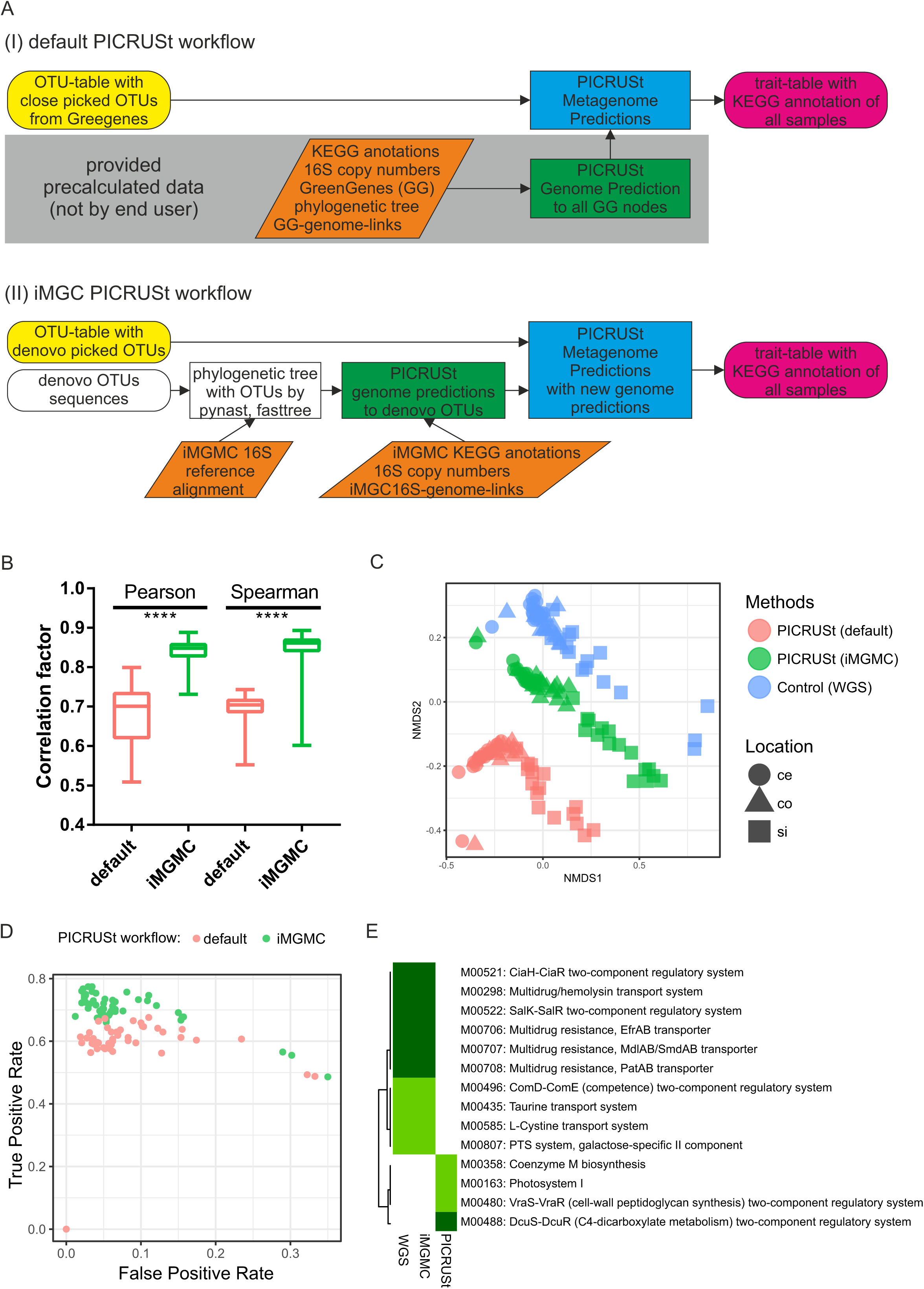
Mouse gut microbiota optimized PICRUStFiMGMC model. (A) The different PICRUSt workflows used in this study: (I) Default workflow for end-user starting from close reference picked OTUs against the GreenGenes database relying on functional metagenome prediction using precalculated genome predictions files (II) Novel PICRUSt workflow starting from *denovo* picked OTUs and using MAGs with 16S rRNA gene links to create ecosystem-specific functional metagenome predictions. (B-E) For comparison of PICRUSt-KEGG-Ortholog (KO) profiles generated using PICRUST-default and PICRUSt-iMGMC from 16S rRNA gene amplicon sequencing to real KO profiles determined by shotgun metagenome sequencing (WGS) samples from different anatomical locations (n=50) were analyzed. (B) Correlation between KO profiles of metagenomes determined by WGS and PICRUST-default (red) or by WGS and PICRUSt-iMGMC (green) using Pearson and Spearman correlation coefficients. ****: p<0.0001 (two-tailed t-test). (C) Comparison of KO profiles generated using PICRUST-default (red), PICRUSt-iMGMC (green) and WGS (blue) from different anatomical locations. Non-metric multidimensional scaling (NMDS) was performed to visualize similarities. (D) False positive rates and true positive rates were obtained by comparing the PICRUSt-default (red) and PICRUSt-iMGMC (green) KEGG Module predictions against WGS results. The true positive rate reflects the fraction of KEGG Modules commonly predicted by both WGS and PICRUSt default/PICRUSt-iMGMC and the false positive rate reflects the fraction of KEGG Modules that are predicted by PICRUSt default/PICRUSt-IMGMC, but were completely absent in WGS data. (E) KEGG module predictions that differ between PICRUSt-default and PICRUSt-iMGMC predictions. KEGG Module prediction by PICRUSt-default and PICRUSt-iMGMC was compared against WGS for all samples and significant differences in completeness were identified using a Wilcoxon test (FDR-corrected). The heatmap displays select KEGG Modules with highly similar completeness between PICRUSt-iMGMC and WGS, but divergent completeness between PICRUSt-default and WGS (see methods for details).

### Multi-scale taxonomic assignment of gene entries based on metagenomic reconstruction enhances taxonomic resolution in iMGMC

Gene catalogs have foremost been employed to generate functional profiles from short read metagenome surveys of communities. To assess the performance of iMGMC in this respect, we performed read-mapping of sequencing data from three external studies, which were not included in the construction of neither iMGMC nor MGCv1, to both catalogs^24–26^. This revealed an increased number of reads (up to 36%) mapping to the iMGMC, supporting the utility of this new catalog (Figure S8).

The taxonomic assignment of entries in classical gene catalogs, specifically after sample-specific assembly and clustering of ORFs by similarity, i.e. 95% identity at 90% coverage in the MGCv1^4^, is limited by the ability of algorithms to predict the taxonomic placement based on relatively short ORFs, which has a limited robustness^27^. Taking advantage of the clustering free approach, we annotated each iMGMC entry using the taxonomic information obtained from the respective gene and contig as well as from the bin ^28^ and the connected MAG/16S rRNA gene sequences, whenever available (Figure 1D). As a result of using longer contigs rather than short ORFs sequences, the relative taxonomic assignment rate improved between 28 and 1,021% at different taxonomic levels (Figure 1D). Notably, many entries were still not assigned to high taxonomic ranks with high confidence, since these approaches are reference-based, and are hampered by the presence of novel and unclassified taxa. Using the MAGs of the iMGMC resource, we could assign up to 40% of mapped reads of three external datasets to MAGs (Figure S8), facilitating the identification of specific bacterial taxa, allowing improved functional analysis by providing information of the genomic context of genes, or of bacterial interaction networks identified by covarying abundances across samples. For instance, the analysis of previously generated shotgun metagenomic data from mice subjected to different experimental diets allowed the retrospective identification of MAG networks rather than gene clusters that show conserved changes in their relative abundance induced by these diets (Figure S9). Hence, future users will be able to utilize in parallel taxonomic information for each gene catalog entry, ranging from well-established methods with lower resolution to innovative methods with enhanced resolution.

### ProviderFspecific diversification of the mouse microbiota

Recent studies have demonstrated that the composition of murine microbiomes varies between different providers, mostly via 16S rRNA amplicon sequence analysis ^29^. However, to which degree laboratory mice share a conserved set of microbes is not known. The presence of a core set of bacteria, based on the detection of 26 CAGs in >95% of mice, was proposed previously ^4^. We analyzed the relative abundance of each individual MAG in all samples by remapping all reads from each library to the MAGs, followed by conversion of mapped read counts into relative abundances (see methods for details). Strikingly, this analysis revealed that each mouse line featured a unique combination of MAGsp even mice from different barriers of the same commercial vendor differed (Figure 2). This resulted in substantial differences in the functional potential of the microbiome within each mouse line (Figure S5D, Table S3). Hence, we next quantitatively assessed the distribution of MAGs by determining their prevalence and relative abundance within each provider. Around 10% of MAGs (70/660) were shared by at least half of the providers (> 0.1% relative abundance in at least one individual sample per provider) (Figure 4A). The most prevalent MAG, matching to *Lactobacillus murinus ASF361* (ANI =97%), was detected in almost all providers (20/21). Notably, three additional members of the Altered Schaedler Flora (ASF) community, which has been studied as mouse gut model community in the past, as well as only four other previously sequenced bacteria were found in at least 50% of providers, while the remaining 62 (=88%) represent uncultured bacteria. We next analyzed the MAGs shared by at least two thirds of the providers (n=21 MAGs) from which most belonged taxonomically to the *Firmicutes* (n=18), two belonged to the *Bacteroidales* S24-7 group (phylum *Bacteroidetes*, proposed family *Muribaculaceae*) and one was identical to *Mucispirillum schaedleri* (phylum *Deferribacteres*) (Figure 4B). Strikingly, the relative abundance of these MAGs revealed large differences between providers (up to 100-fold) suggesting that their respective abundance within each community is strongly influenced by environmental factors. Taking advantage of the link between MAG and 16S rRNA gene sequences, we assessed the global prevalence and relative abundance of the corresponding 16S rRNA gene sequences across all 16S rRNA amplicon datasets deposited in the SRA using the recently established IMNGS database (Figure 4C)^30^. This search revealed that the most prevalent MAG in our study, *Lactobacillus murinus,* is present in 36% of all samples derived from the mouse gut (n=9,496), while being largely absent from the human gut and only detectable in 1.4% of rat gut microbiota samples (1.4% positive) (Table S4). To assess whether the newly reconstructed 16S rRNA gene sequences represented taxa commonly found in mice, we employed IMNGS and queried all 1,323 16S rRNA gene sequences to assess their relative abundance in SRA samples derived from diverse ecosystems (Figure 4D and E). A prevalence of 1% (threshold relative abundance: 0.1%) within at least one of the ecosystems was determined for 739 rRNA gene sequences from which 569 were enriched in the mouse gut, mouse skin, rat gut or human gut. Of these 44% were most prevalent in the mouse gut, with an additional 6% being shared with the mouse skin. Other sequences were shared with the rat microbiome (12%) and the human gut microbiome (7%) (Figure 4E). In summary, our large-scale analysis revealed the presence of specific bacteria commonly found in mouse lines but no other gut microbiomes, yet, also a high species-level variability within the murine gut microbiome, which impacts the functional repertoire of the microbiome and potentially thereby the outcome of *in vivo* experiments.

**Figure 4:**
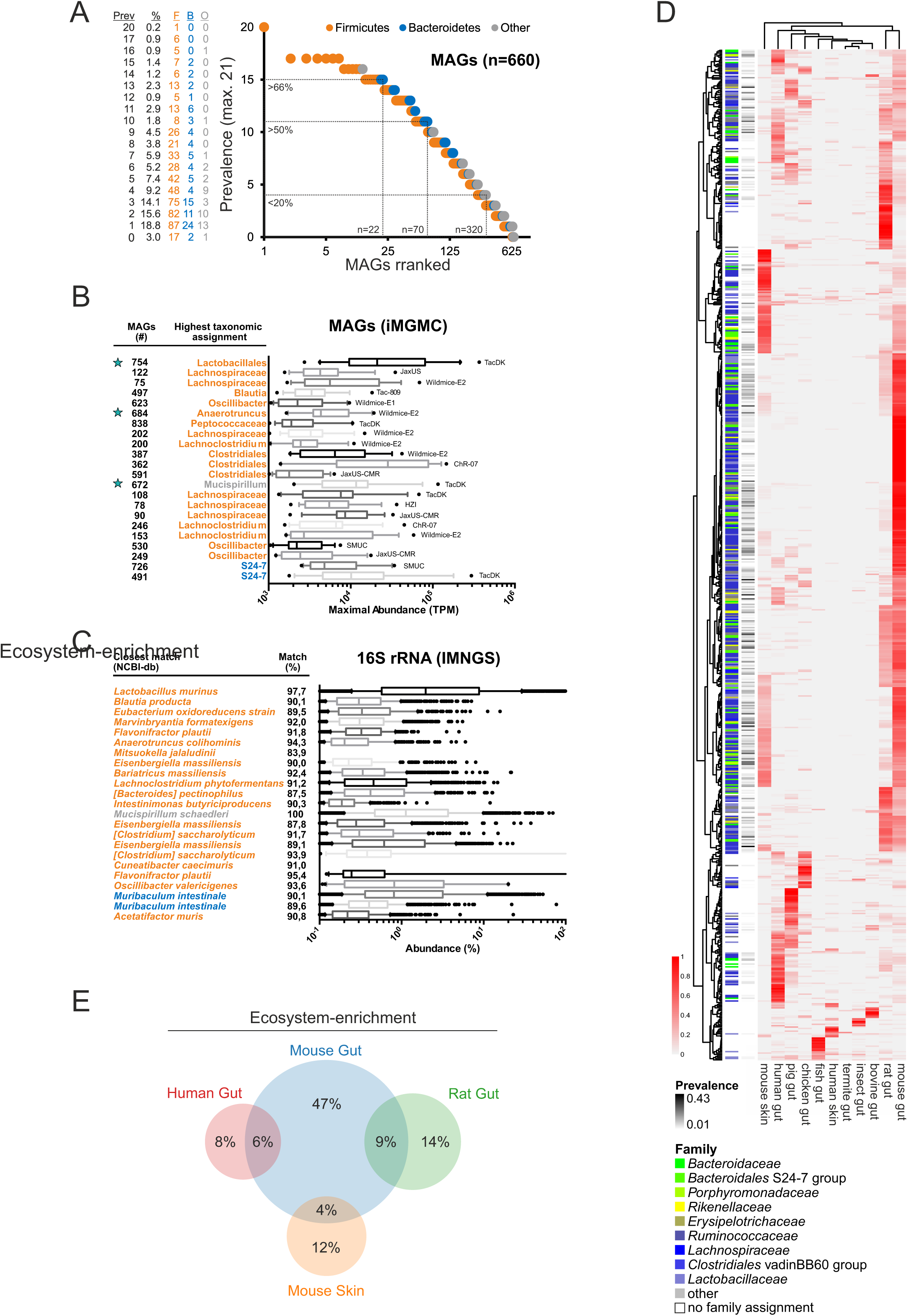
Identification of MAGs shared between laboratory mice. (A) Prevalence of MAGs (n=660) in samples from 21 mouse providers. MAGs were considered present in a provider if its relative abundance reached at least 0.1% in one sample of the provider. Numberson the left indicate the fraction (%) and taxonomic grouping (F: Firmicutes, B: Bacteroides, O: Other phyla) of MAGs with an indicated prevalence (Prev). In the right panel MAGs were ranked by prevalence and dashed lines indicate number of MAGs present in >66%, >50% and >20% of providers, respectively.. (B) Comparison of maximal abundance between providers for each MAG (n=22) present in at least 2/3 of providers. For each MAG, the bin number, the highest taxonomic assignment based on the manually curated phylogenetic tree and the provider with the highest abundance is listed. Stars indicate MAGs with matches in NCBI RefSeq. (C) Comparison of the relative abundance of 16S rRNA gene sequences linked to MAGs in the IMNGS database. For each 16S rRNA gene, the closest named relative 16S rRNA gene sequence was determined and blasted to the NCBI-16S rRNA gene database. Color of dots and names indicate their taxonomic association to different phyla (F: Firmicutes, B: Bacteroidetes, O: other phyla) (D and E) IMNGS was used to determine the prevalence for iMGMC 16S rRNA gene sequences (n=1,323) in distinct hosts and ecosystems. Of these 1,113 reached at least a prevalence threshold of 1% prevalence within one of the evaluated environment (0.1% sample-depth cutoff of presence). Resulting sequences (n=1,113) were filtered further to have at least >1% relative mean abundance in at least one environment. (D) Heatmap displaying the mean relative abundance within an ecosystem (row normalized) of those 16S rRNA gene sequences which have at least >1% relative mean abundance in at least one environment (n=739). (E) Venn diagram visualizing the distribution of 16S rRNA gene sequences subsampled to be enriched (>50% relative abundance normalized over the ecosystems in Figure 4D) in mouse gut, mouse skin, rat gut and human gut microbiome (n = 569). Numbers indicate fraction of 16S rRNA gene sequences enriched or shared between indicated ecosystems.

## Discussion

Short read-based sequencing studies of microbial ecosystems require suitable reference databases for maximal resolution of taxonomic and functional assignments. Gene catalogs and 16S rRNA gene databases commonly represent separate references for shotgun metagenome and 16S rRNA amplicon sequencing analyses, respectively. To overcome the separation between these types of databases, a novel framework that can serve as i) a valuable resource for the most utilized experimental model for microbiome research, the mouse gut microbiota, and ii) a blue print to generate integrated gene catalogs for less characterized microbial ecosystems was developed.

For the establishment of the integrated gene catalog, methods identified to yield optimal results by the CAMI challenge^27^, e.g. for assembly of MAGs or binning when dealing with large datasets, were utilized and complemented with a novel approach linking MAGs and 16S rRNA sequences. The “All-in-One” assembly resulted for the mouse gut microbiome in the reconstruction of a large number of high-quality MAGs, including low abundant community members, representing bacteria that were neither cultured or identified in other high-throughput sequencing studies^17^. Strikingly, for both the mouse and pig gut microbiome, more than 87% of MAGs fell into this category. The *Clostridiales*-vadinBB660 or *Mollicutes* RF9 groups, which were so far only known from 16S rRNA gene sequencing, are examples of functionally distinct and underexplored bacteria frequently occurring in mouse gut microbiomes. Preliminary analysis of assemblies of large datasets from the human gut microbiome suggest that the developed approach also identifies hundreds of novel MAGs (approximately 30% of assembled MAGs), demonstrating the power of this approach even for better characterized ecosystems.

Another utility of the integrated gene catalog is the availability of linked MAG-16S rRNA gene pairs, which enables the incorporation of data from large 16S rRNA gene databases such as the IMNGS database encompassing 168,573 short-read datasets (build 1711) thereby allowing large-scale screening for identified MAGs, such as the evaluation of a core microbiome in the mouse gut. The MAG-16S rRNA gene pairs also enabled the development of an ecosystem-optimized version of PICRUSt, which produced gene profiles more closely resembling WGS data. We anticipate this to be widely adapted to predict metagenome profiles based on 16S rRNA amplicon sequencing data and suggest that ecosystem-optimized versions of PICRUSt will be valuable resources.

Altogether, the clustering-free construction of gene catalogs together with the reconstruction of a large number of almost complete MAGs through an improved assembly strategy as well as linking to 16S rRNA gene sequences provide a highly-integrated resource for sequencing-based work and will enable future studies to explore the taxonomy, functionality and community structure of the mouse gut and other ecosystems in more depth.

## Acknowledgements

TS was funded by the Helmholtz Association (VH-NG-933), by the Deutsche Forschungsgemeinschaft (DFG, German Research Foundation, STR-1343/1 and STR-1343/2) and the European Union (StG337251).

JFB was funded by the DFG under Germany’s Excellence Strategy – EXC 22167-390884018 and by the DFG Collaborative Research Center (CRC) 1182 “Origin and Function of Metaorganisms”.

TC received funding from the DFG (CL481/2-1).

## References

1. Kamada, N., Seo, S.\U., Chen, G. Y. & Núñez, G. Role of the gut microbiota in immunity and inflammatory disease. Nat. Rev. Immunol. 13, 321–35 (2013).

2. Li, J. et al. An integrated catalog of reference genes in the human gut microbiome. Nat. Biotechnol. 32, 834–841 (2014).

3. Sunagawa, S. et al. Structure and function of the global ocean microbiome. Science (80H.). 348, 1261359–1261359 (2015).

4. Xiao, L. et al. A catalog of the mouse gut metagenome. Nat. Biotechnol. 33, 1103–1108 (2015).

5. Xiao, L. et al. A reference gene catalogue of the pig gut microbiome. Nat. Microbiol. 1, 16161 (2016).

6. Lagkouvardos, I. et al. The Mouse Intestinal Bacterial Collection (miBC) provides host\specific insight into cultured diversity and functional potential of the gut microbiota. Nat. Microbiol. 1, 16131 (2016).

7. Li, D. et al. MEGAHIT v1.0: A fast and scalable metagenome assembler driven by advanced methodologies and community practices. Methods 102, 3–11 (2016).

8. Zhu, W., Lomsadze, A. & Borodovsky, M. Ab initio gene identification in metagenomic sequences. Nucleic Acids Res. 38, e132–e132 (2010).

9. Fu, L., Niu, B., Zhu, Z., Wu, S. & Li, W. CD\HIT: accelerated for clustering the next\generation sequencing data. Bioinformatics 28, 3150–3152 (2012).

10. Kang, D. D., Froula, J., Egan, R. & Wang, Z. MetaBAT, an efficient tool for accurately reconstructing single genomes from complex microbial communities. PeerJ 3, e1165 (2015).

11. Parks, D. H., Imelfort, M., Skennerton, C. T., Hugenholtz, P. & Tyson, G. W. CheckM: assessing the quality of microbial genomes recovered from isolates, single cells, and metagenomes. Genome Res. 25, 1043–1055 (2015).

12. Miller, C. S., Baker, B. J., Thomas, B. C., Singer, S. W. & Banfield, J. F. EMIRGE: reconstruction of full\length ribosomal genes from microbial community short read sequencing data. Genome Biol. 12, R44 (2011).

13. Zeng, F., Wang, Z., Wang, Y., Zhou, J. & Chen, T. Large\scale 16S gene assembly using metagenomics shotgun sequences. Bioinformatics 33, 1447–1456 (2017).

14. Delmont, T. O. & Eren, A. M. Identifying contamination with advanced visualization and analysis practices: metagenomic approaches for eukaryotic genome assemblies. PeerJ 4, e1839 (2016).

15. Mikheenko, A., Saveliev, V. & Gurevich, A. MetaQUAST: evaluation of metagenome assemblies. Bioinformatics 32, 1088–1090 (2016).

16. Alneberg, J. et al. Binning metagenomic contigs by coverage and composition. Nat. Methods 11, 1144–1146 (2014).

17. Parks, D. H. et al. Recovery of nearly 8,000 metagenome\assembled genomes substantially expands the tree of life. Nat. Microbiol. 2, 1533–1542 (2017).

18. Parks, D. H. et al. A standardized bacterial taxonomy based on genome phylogeny substantially revises the tree of life. Nat. Biotechnol. 36, 996–1004 (2018).

19. Clavel, T., Lagkouvardos, I., Blaut, M. & Stecher, B. The mouse gut microbiome revisited: From complex diversity to model ecosystems. Int. J. Med. Microbiol. 306, 316–327 (2016).

20. Karst, S. M. et al. Retrieval of a million high\quality, full\length microbial 16S and 18S rRNA gene sequences without primer bias. Nat. Biotechnol. 36, 190–195 (2018).

21. Langille, M. G. I. et al. Predictive functional profiling of microbial communities using 16S rRNA marker gene sequences. Nat. Biotechnol. 31, 814–821 (2013).

22. Aßhauer, K. P., Wemheuer, B., Daniel, R. & Meinicke, P. Tax4Fun: predicting functional profiles from metagenomic 16S rRNA data: Fig. 1. Bioinformatics 31, 2882–2884 (2015).

23. Buchfink, B., Xie, C. & Huson, D. H. Fast and sensitive protein alignment using DIAMOND. Nat. Methods 12, 59–60 (2015).

24. Suez, J. et al. Artificial sweeteners induce glucose intolerance by altering the gut microbiota. Nature 514, 181–186 (2014).

25. Everard, A. et al. Microbiome of prebiotic\treated mice reveals novel targets involved in host response during obesity. ISME J. 8, 2116–2130 (2014).

26. Levy, M. et al. Microbiota\Modulated Metabolites Shape the Intestinal Microenvironment by Regulating NLRP6 Inflammasome Signaling. Cell 163, 1428–1443 (2015).

27. Sczyrba, A. et al. Critical Assessment of Metagenome Interpretation—a benchmark of metagenomics software. Nat. Methods 14, 1063–1071 (2017).

28. Cambuy, Diego D, Coutinho, Felipe H, Dutilh, B. E. Contig annotation tool CAT robustly classifies assembled metagenomic contigs and long sequences. BioRxiv 072868 (2016). doi:10.1101/072868

29. Rausch, P. et al. Analysis of factors contributing to variation in the C57BL/6J fecal microbiota across German animal facilities. Int. J. Med. Microbiol. 306, (2016).

30. Lagkouvardos, I. et al. IMNGS: A comprehensive open resource of processed 16S rRNA microbial profiles for ecology and diversity studies. Sci. Rep. 6, 33721 (2016).

