## Supplementary Information for "An integrated metagenome catalog reveals novel insights into the murine gut microbiome"

Affiliations:
1: Department of Microbial Immune Regulation, Helmholtz Centre for Infection Research, Braunschweig, Germany.
2: ZIEL Institute for Food and Health, Technical University of Munich, Freising, Germany
3: Max Planck Institute for Evolutionary Biology, Plön, Germany.
4: Institute for Experimental Medicine, Kiel University, Kiel, Germany.
5: Functional Microbiome Research Group, Institute of Medical Microbiology, RWTH University Hospital, Aachen, Germany
6: Center for Biotechnology, Bielefeld University, Bielefeld, Germany.
7: Department of Computational Biology of Infection Research, Helmholtz Centre for Infection Research, Braunschweig, Germany.
8: Braunschweig Integrated Centre of Systems Biology, Braunschweig, Germany
9: Hanover Medical School, Hannover, Germany
10: RESIST, Cluster of Excellence 2155, Hanover Medical School, Hanover, Germany

### Supplementary Figure Legends

**Figure S01: Flowchart of methodology to link MAGs to 16S rRNA gene sequences by combining mapping-based and statistical approaches**

Flowchart of the methodology to link MAGs to 16S rRNA gene sequences by combining mapping-based and statistical approaches. Metagenomic bins and 16S rRNA gene sequences were assembled in an “all-in-one”-approach from 298 metagenomic libraries from the murine gut microbiota using MEGAHIT/MetaBat and RAMBL, respectively. A total of 1,462 bins (>200 kbp) and 1,323 reconstructed 16S rRNA gene sequences were used as input for the linkage of bins to 16S rRNA gene sequences. The methodology relies on three levels of analysis to generate integrated scores reflecting linkage confidence: (I) Search for 16S rRNA gene sequences in the bins using Blastn [1]; (II) Identify read-pairs that contain a 16S rRNA gene sequence in one read and a sequence from a bin in the other thereby enabling a link between bins to reconstructed 16S rRNA gene sequences (“scaffolding”); and (III) Determine co-abundance profiles of bins and 16S rRNA gene sequences through adaptation of a previously described correlation-based approach [2]. All results were transformed and combined into an integrated scoring scheme to identify the “best-hit” for bin/16S rRNA gene sequence pairs.

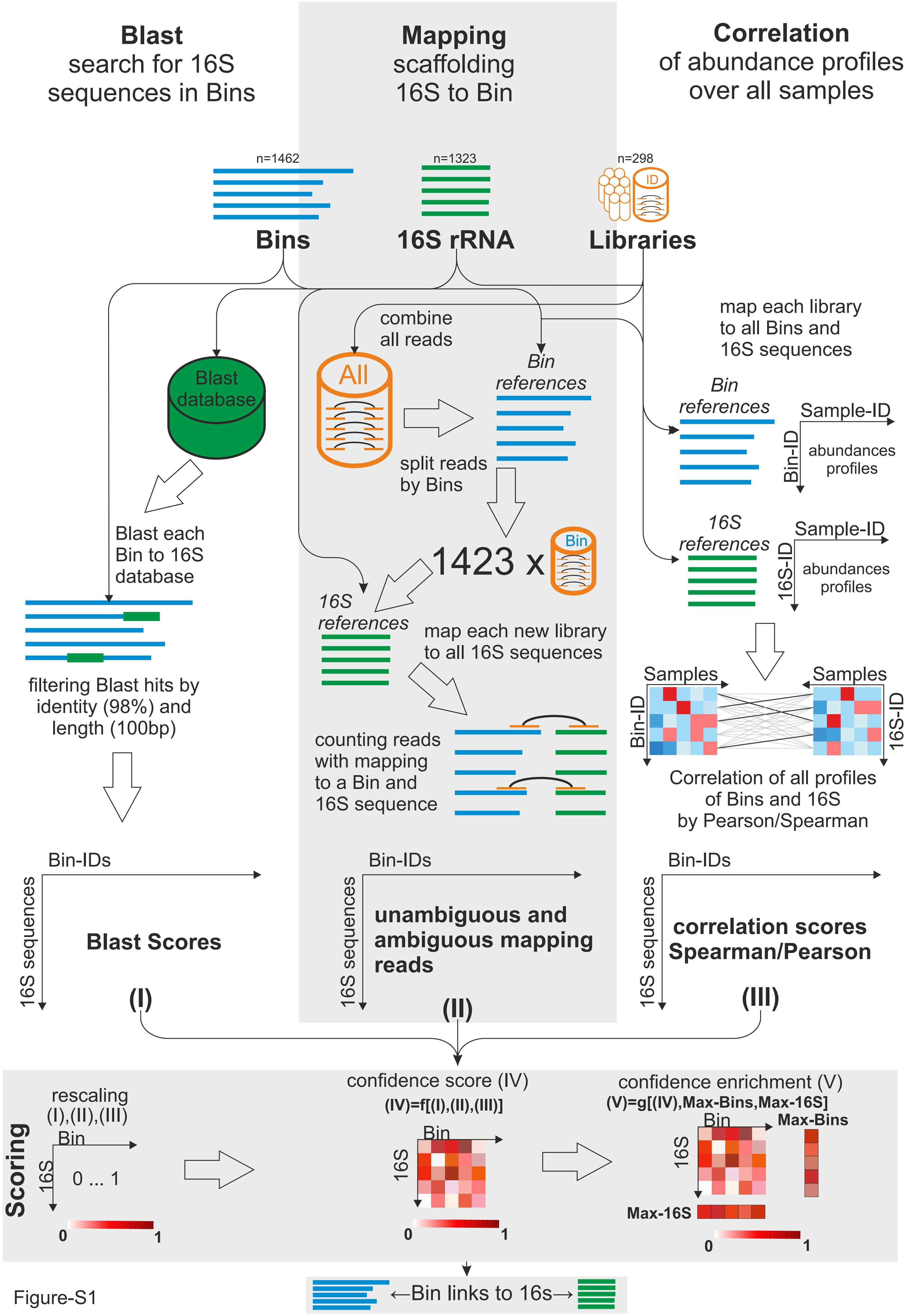

Blast
search for 16S sequences in Bins
Mapping
scaffolding 16S to Bin
Correlation
of abundance profiles over all samples
n=1462
n=1323
n=298
ID
Bins
16S rRNA
Libraries
combine all reads
All
Blast database
Bin references
split reads by Bins
map each library to all Bins and 16S sequences
Bin references
Sample-ID
Bin-ID
abundances profiles
16S references
Sample-ID
16S-ID
abundances profiles
Blast each Bin to 16S database
1423 x
Bin
16S references
map each new library to all 16S sequences
filtering Blast hits by identity (98%) and length (100bp)
counting reads with mapping to a Bin and 16S sequence
Samples
Samples
Bin-ID
16S-ID
Correlation of all profiles of Bins and 16S by Pearson/Spearman
Bin-IDs
16S sequences
Blast Scores
(I)
Bin-IDs
16S sequences
unambiguous and ambiguous mapping reads
(II)
Bin-IDs
16S sequences
correlation scores Spearman/Pearson
(III)
Scoring
rescaling (I),(II),(III)
Bin
16S
0 ... 1
0
1
confidence score (IV)
(IV)=f[(I),(II),(III)]
Bin
16S
0
1
confidence enrichment (V)
(V)=g[(IV),Max-Bins,Max-16S]
Bin
Max-Bins
16S
Max-16S
0
1
←Bin links to 16s→


Figure-S1

**Figure S02: Evaluation of the binning efficiency by using know NCBI reference genomes**

(A) Pipeline to identify closely related NCBI Ref genomes present in the assembly. Identified genomes were subsequently used for binning evaluation.

(B) Stacked bar plots indicated the genome coverage of reference genome in assembly. The complete bar indicates the overall genome coverage within contigs. The part of the genome which is mapped to bins (green: largest bin, yellow, second largest bin, orange: rest of bins) or to unbinned contigs (red) is indicated. The number in the stacked bar represents the MAG number (see Table S2).

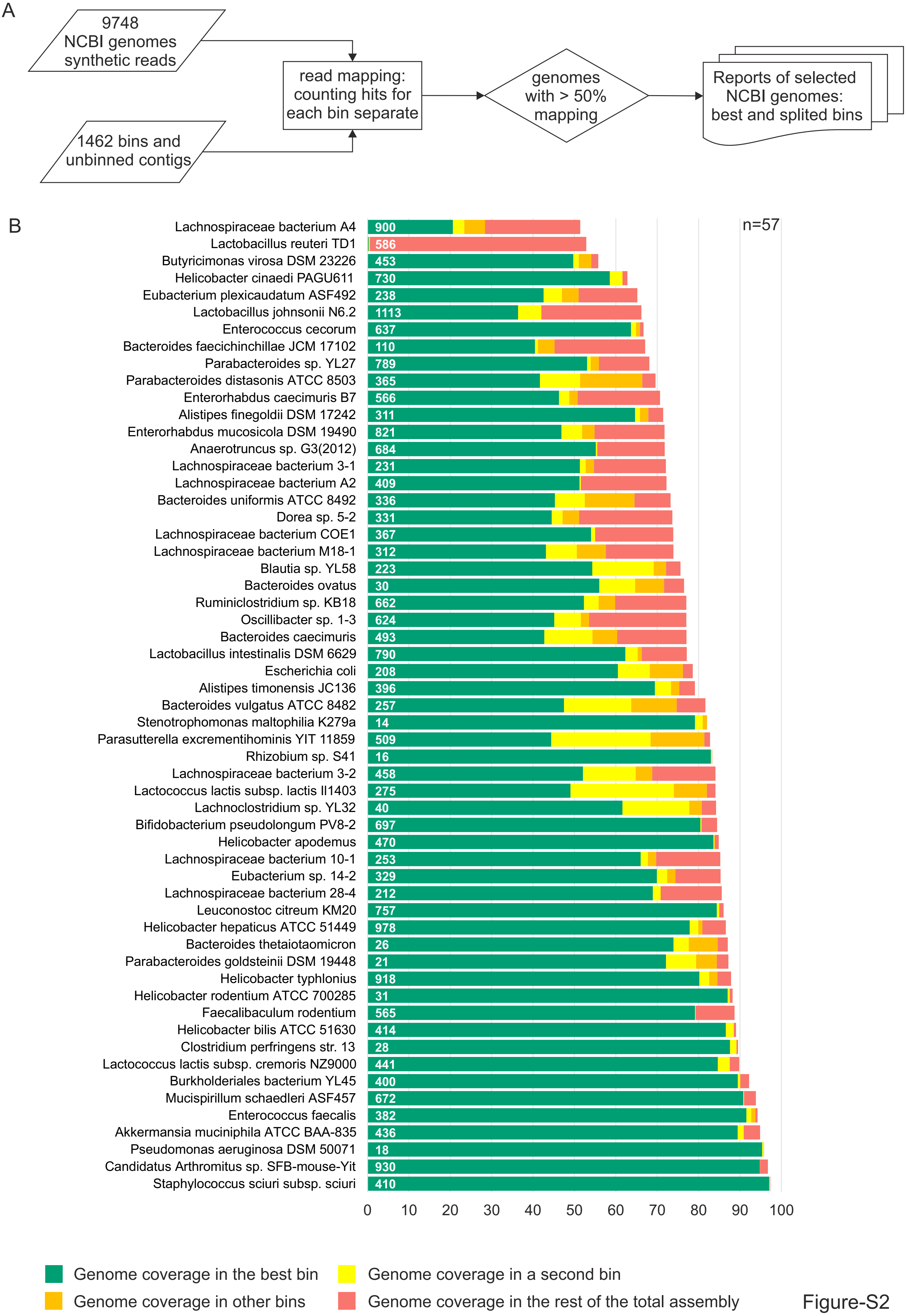
A
9748
NCBI genomes
synthetic reads
1462 bins and
unbinned contigs
read mapping:
counting hits for
each bin separate
genomes
with > 50%
mapping
Reports of selected
NCBI genomes:
best and splited bins
B
n=57
Lachnospiraceae bacterium A4 900
Lactobacillus reuteri TD1 586
Butyricimonas virosa DSM 23226 453
Helicobacter cinaedi PAGU611 730
Eubacterium plexicaudatum ASF492 238
Lactobacillus johnsonii N6.2 1113
Enterococcus cecorum 637
Bacteroides faecichinchillae JCM 17102 110
Parabacteroides sp. YL27 789
Parabacteroides distasonis ATCC 8503 365
Enterorhabdus caecimuris B7 566
Alistipes finegoldii DSM 17242 311
Enterorhabdus mucosicola DSM 19490 821
Anaerotruncus sp. G3(2012) 684
Lachnospiraceae bacterium 3-1 231
Lachnospiraceae bacterium A2 409
Bacteroides uniformis ATCC 8492 336
Dorea sp. 5-2 331
Lachnospiraceae bacterium COE1 367
Lachnospiraceae bacterium M18-1 312
Blautia sp. YL58 223
Bacteroides ovatus 30
Ruminiclostridium sp. KB18 662
Oscillibacter sp. 1-3 624
Bacteroides caecimuris 493
Lactobacillus intestinalis DSM 6629 790
Escherichia coli 208
Alistipes timonensis JC136 396
Bacteroides vulgatus ATCC 8482 257
Stenotrophomonas maltophilia K279a 14
Parasutterella excrementihominis YIT 11859 509
Rhizobium sp. S41 16
Lachnospiraceae bacterium 3-2 458
Lactococcus lactis subsp. lactis Il1403 275
Lachnoclostridium sp. YL32 40
Bifidobacterium pseudolongum PV8-2 697
Helicobacter apodemus 470
Lachnospiraceae bacterium 10-1 253
Eubacterium sp. 14-2 329
Lachnospiraceae bacterium 28-4 212
Leuconostoc citreum KM20 757
Helicobacter hepaticus ATCC 51449 978
Bacteroides thetaiotaomicron 26
Parabacteroides goldsteinii DSM 19448 21
Helicobacter typhlonius 918
Helicobacter rodentium ATCC 700285 31
Faecalibaculum rodentium 565
Helicobacter bilis ATCC 51630 414
Clostridium perfringens str. 13 28
Lactococcus lactis subsp. cremoris NZ9000 441
Burkholderiales bacterium YL45 400
Mucispirillum schaedleri ASF457 672
Enterococcus faecalis 382
Akkermansia muciniphila ATCC BAA-835 436
Pseudomonas aeruginosa DSM 50071 18
Candidatus Arthromitus sp. SFB-mouse-Yit 930
Staphylococcus sciuri subsp. sciuri 410
0 10 20 30 40 50 60 70 80 90 100
Genome coverage in the best bin
Genome coverage in a second bin
Genome coverage in other bins
Genome coverage in the rest of the total assembly


Figure-S2

**Figure S03: Evaluation of linking reconstructed 16S rRNA gene sequences to MAG/bins**

(A) NCBI reference genomes were to evaluate the links between MAG/16S rRNA gene links (n= 484). Specifically, those reference genomes, which were contained within MAGs, were utilized to compare their reference 16S rRNA gene sequence against the 1,323 reconstructed 16S rRNA gene sequences and then against the linked 16S rRNA gene sequence of the respective MAG. Numbers (II-VI) indicate the comparison against different scoring parameters in panel B.

(B) (I) The colored bars represent taxonomic agreement at different levels of linked MAGs and reconstructed 16S rRNA gene sequences (red = different; green = overlapping taxonomic assignment). Stars indicate the best possible matching of reconstructed 16S rRNA gene sequence, based on matched NCBI genome and their corresponding 16S rRNA gene sequence obtained by the scoring scheme described above. (II) Identity (%) of the 16S rRNA gene sequence inferred from the NCBI-genome to the reconstructed 16S rRNA gene sequence of highest identity via BlastN. (III) Blast scoring of 16S rRNA gene sequences present in the MAG/bin. (IV) Number of reads mapped to 16S rRNA gene sequences and to the selected MAG/bin in the scaffolding step. (V) Correlation score of abundances profiles of 16S rRNA gene and MAGs sequences via mapping rates. (VI) Integrated score for linking 16S rRNA genes and the MAG/bin.

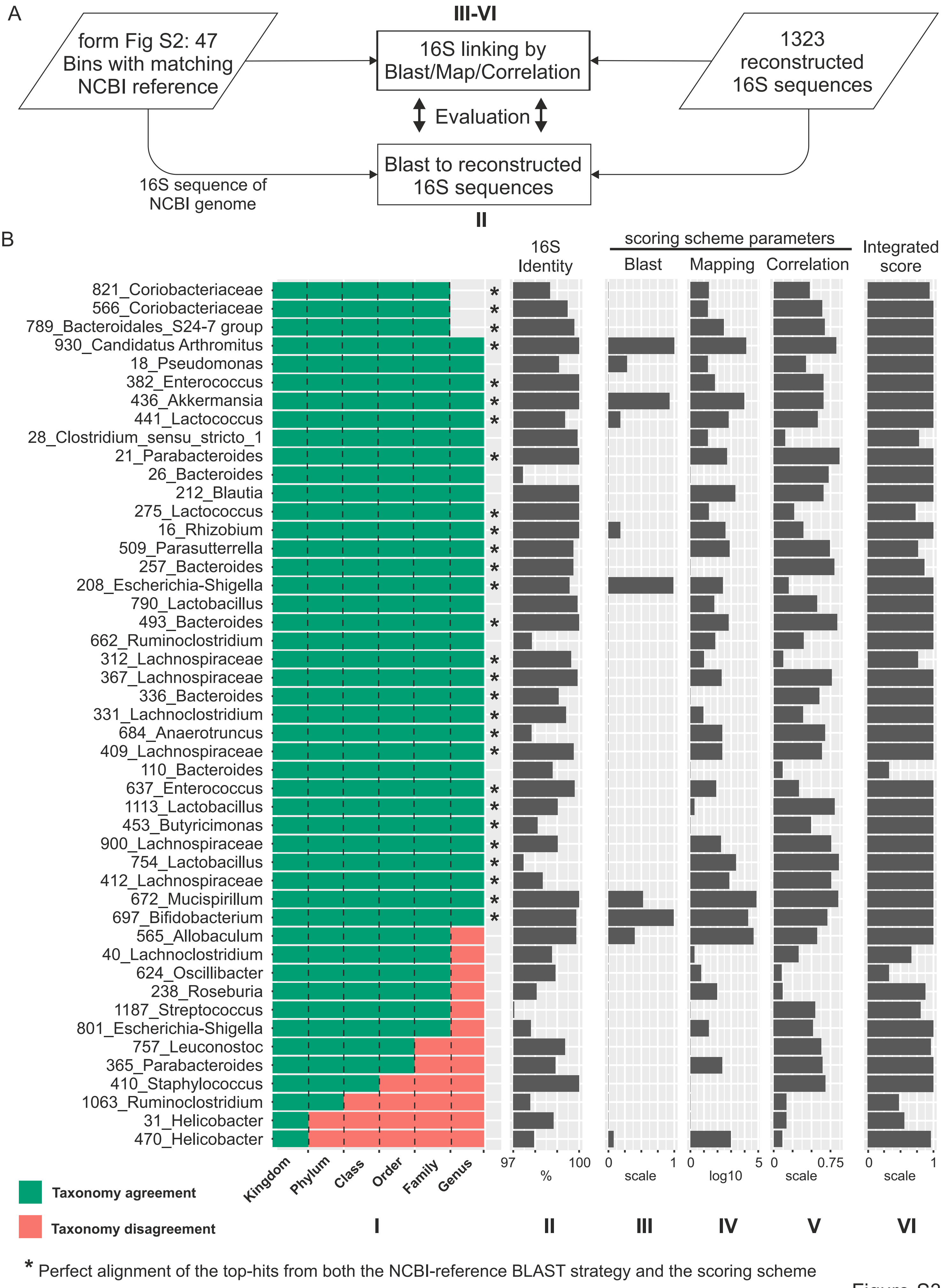
A
III-VI
form Fig S2: 47 Bins with matching NCBI reference
16S linking by Blast/Map/Correlation
1323 reconstructed 16S sequences
Evaluation
Blast to reconstructed 16S sequences
16S sequence of NCBI genome
II
B
16S Identity
scoring scheme parameters
Blast
Mapping
Correlation
Integrated score
821_Coriobacteriaceae
566_Coriobacteriaceae
789_Bacteroidales_S24-7 group
930_Candidatus Arthromitus
18_Pseudomonas
382_Enterococcus
436_Akkermansia
441_Lactococcus
28_Clostridium_sensu_stricto_1
21_Parabacteroides
26_Bacteroides
212_Blautia
275_Lactococcus
16_Rhizobium
509_Parasutterrella
257_Bacteroides
208_Escherichia-Shigella
790_Lactobacillus
493_Bacteroides
662_Ruminoclostridium
312_Lachnospiraceae
367_Lachnospiraceae
336_Bacteroides
331_Lachnoclostridium
684_Anaerotruncus
409_Lachnospiraceae
110_Bacteroides
637_Enterococcus
1113_Lactobacillus
453_Butyricimonas
900_Lachnospiraceae
754_Lactobacillus
412_Lachnospiraceae
672_Mucispirillum
697_Bifidobacterium
565_Allobaculum
40_Lachnoclostridium
624_Oscillibacter
238_Roseburia
1187_Streptococcus
801_Escherichia-Shigella
757_Leuconostoc
365_Parabacteroides
410_Staphylococcus
1063_Ruminoclostridium
31_Helicobacter
470_Helicobacter
Kingdom
Phylum
Class
Order
Family
Genus
97
100
%
0
1
scale
0
5
log10
0
0.75
scale
0
1
scale
Taxonomy agreement
Taxonomy disagreement
I
II
III
IV
V
VI
* Perfect alignment of the top-hits from both the NCBI-reference BLAST strategy and the scoring scheme

Figure-S3

**Figure S04: Taxonomic diversity of MAGs and reconstructed 16S rRNA gene sequences.**

(A) The sunburst graphs visualize the taxonomic diversity of MAGs (left panel, n=660) and **reconstructed 16S rRNA gene sequences (right panel, n=1,323)**. MAGs/16S RNA gene sequences were grouped according to their lowest taxonomic assignment determined using GTDBTk and SINA, respectively. Each level in the graphs corresponds to a taxonomic level from phylum (inner ring) to genus (outer ring). Letters indicate taxonomic groups with at least 10 MAGs. Numbers in brackets indicate the number of MAGs/16S RNA belonging to the respective taxonomic group.

(B) A phylogentic tree containing the reconstructed 16S rRNA gene sequences. Taxonomic groups are highlighted. The color in the outer ring indicates the presence of a linked MAGs (blue) or CAG (green).

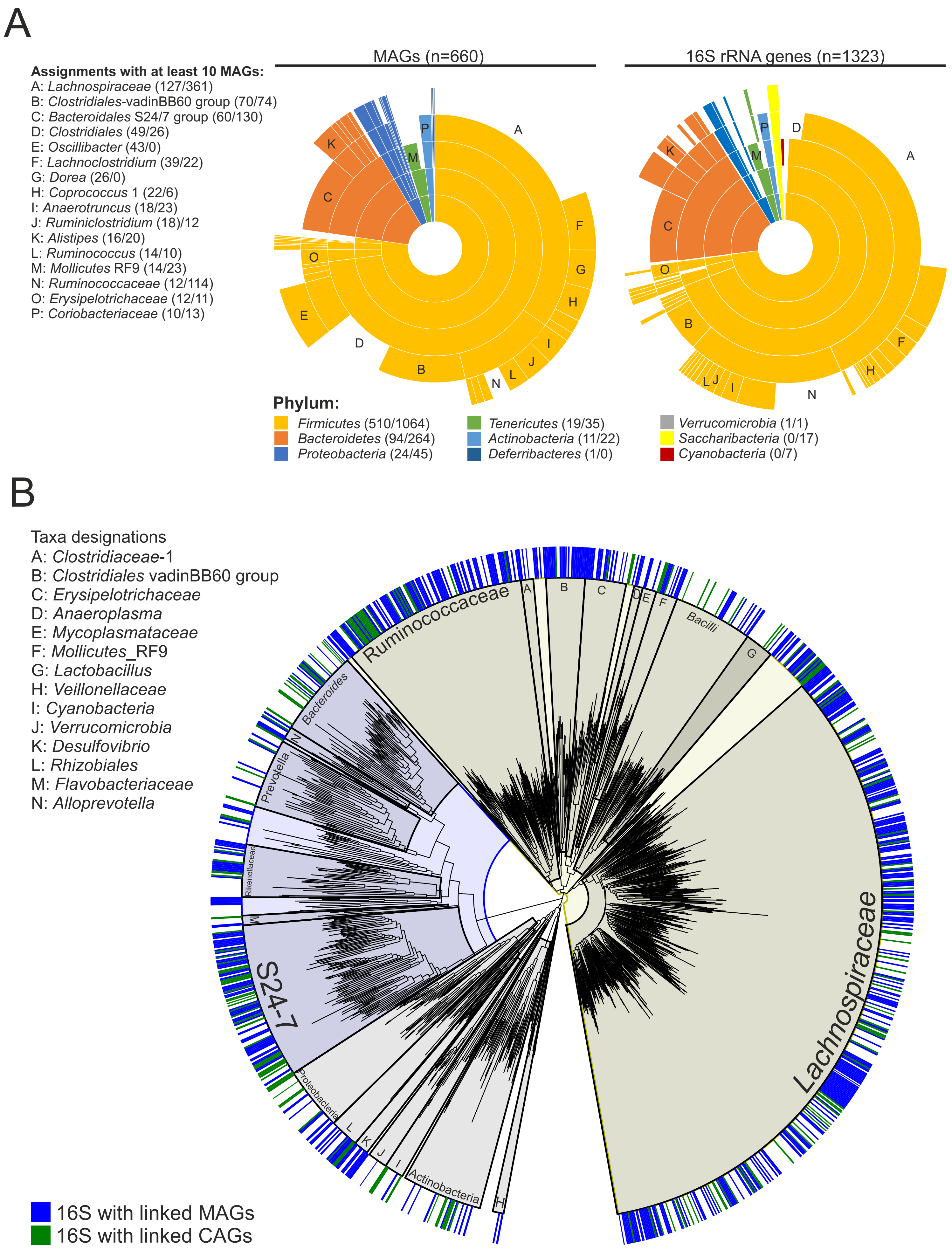
A
Assignments with at least 10 MAGs:
A: Lachnospiraceae (127/361)
B: Clostridiales-vadinBB60 group (70/74)
C: Bacteroidales S24/7 group (60/130)
D: Clostridiales (49/26)
E: Oscillibacter (43/0)
F: Lachnoclostridium (39/22)
G: Dorea (26/0)
H: Coprococcus 1 (22/6)
I: Anaerotruncus (18/23)
J: Ruminiclostridium (18)/12
K: Alistipes (16/20)
L: Ruminococcus (14/10)
M: Mollicutes RF9 (14/23)
N: Ruminococcaceae (12/114)
O: Erysipelotrichaceae (12/11)
P: Coriobacteriaceae (10/13)
MAGs (n=660)
16S rRNA genes (n=1323)
Phylum:
Firmicutes (510/1064)
Bacteroidetes (94/264)
Proteobacteria (24/45)
Tenericutes (19/35)
Actinobacteria (11/22)
Deferribacteres (1/0)
Verrucomicrobia (1/1)
Saccharibacteria (0/17)
Cyanobacteria (0/7)
B
Taxa designations
A: Clostridiaceae-1
B: Clostridiales vadinBB60 group
C: Erysipelotrichaceae
D: Anaeroplasma
E: Mycoplasmataceae
F: Mollicutes_RF9
G: Lactobacillus
H: Veillonellaceae
I: Cyanobacteria
J: Verrucomicrobia
K: Desulfovibrio
L: Rhizobiales
M: Flavobacteriaceae
N: Alloprevotella
Ruminococcaceae
Bacilli
Bacteroides
Prevotella
Rikenellaceae
S24-7
Proteobacteria
Actinobacteria
Lachnospiraceae
16S with linked MAGs
16S with linked CAGs

**Figure S05: Analysis of functional diversity within bacterial members of the mouse gut microbiota and between mouse providers using iMGMC.**

(A-C) Ordination analysis of functional profiles of MAGs contained in iMGMC based on the presence of KEGG orthologs (KO). Comparison of all MAGs (B, n=660) as well as those with taxonomic assignment to the orders Bacteroidales (C, n=94) and Clostridiales (D, n=482). The distances reflect the differences in the functional capabilities of the MAGs according to the presence of KO. Colors represent different taxonomic clusters according to the manually curated phylogenetic MAGs tree (see Figure 2).

(D) To characterize the functional potential of each providers microbiome, individual libraries (n=299) were mapped to the iMGMC. The mapped reads were used to quantify KOs present in each library. This information was translated to KEGG Module completeness scores using KEGGs "Reconstruct Module" function and summarized per provider. The completeness of each KEGG modules was expressed using a color code from dark green (module complete) to white (module absent).

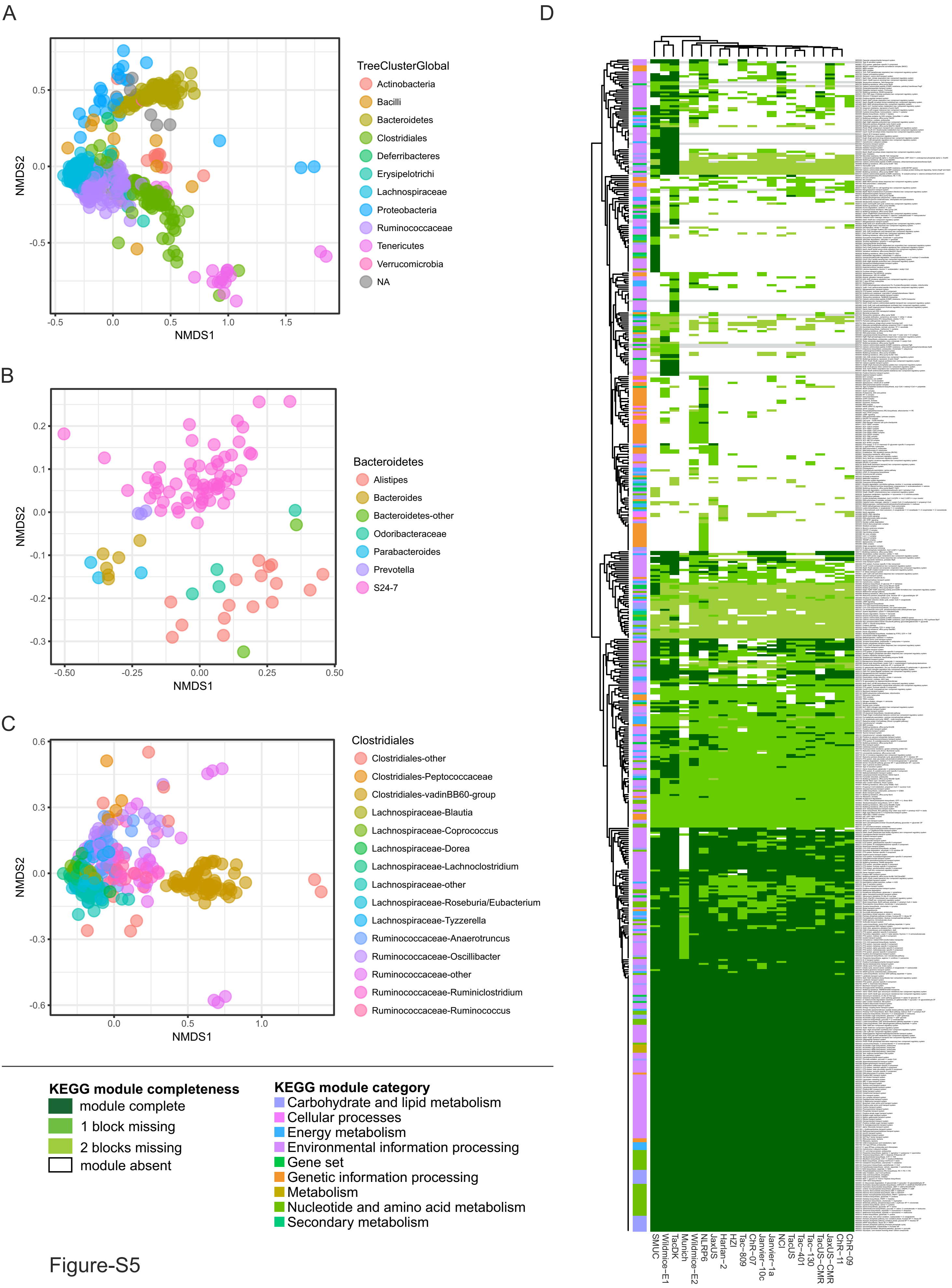
A
B
C
D
TreeClusterGlobal
Actinobacteria
Bacilli
Bacteroidetes
Clostridiales
Deferribacteres
Erysipelotrichi
Lachnospiraceae
Proteobacteria
Ruminococcaceae
Tenericutes
Verrucomicrobia
NA
NMDS1
NMDS2
Bacteroidetes
Alistipes
Bacteroides
Bacteroidetes-other
Odoribacteraceae
Parabacteroides
Prevotella
S24-7
Clostridiales
Clostridiales-other
Clostridiales-Peptococcaceae
Clostridiales-vadinBB60-group
Lachnospiraceae-Blautia
Lachnospiraceae-Coprococcus
Lachnospiraceae-Dorea
Lachnospiraceae-Lachnoclostridium
Lachnospiraceae-other
Lachnospiraceae-Roseburia/Eubacterium
Lachnospiraceae-Tyzzerella
Ruminococcaceae-Anaerotruncus
Ruminococcaceae-Oscillibacter
Ruminococcaceae-other
Ruminococcaceae-Ruminiclostridium
Ruminococcaceae-Ruminococcus
KEGG module completeness
module complete
1 block missing
2 blocks missing
module absent
KEGG module category
Carbohydrate and lipid metabolism
Cellular processes
Energy metabolism
Environmental information processing
Gene set
Genetic information processing
Metabolism
Nucleotide and amino acid metabolism
Secondary metabolism
SMUC
Wildmice−E1
TacDK
Munich
Wildmice−E2
NLRP6
JaxUS
Harlan−2
HZI
Tac−809
ChR−07
Janvier−10c
Janvier−1a
NCI
TacUS
Tac−401
Tac−130
TacUS−CMR
JaxUS−CMR
ChR−11
ChR−09

Figure-S5

**Figure S6: Evaluation of mouse microbiota optimized PICRUSt model built from 484 iMGMC-MAGs with unique 16S rRNA sequences**

Correlation between KO profiles of metagenomes (n=50, fecal samples used to generate the iMGMC) determined by WGS and different PICRUST-workflows using Pearson (A) and Spearman correlation (B), respectively. Correlations factor were determinate for different anatomical locations separately (C and D). Two-tailed paired t-test was performed to analyze the differences, *: $p<0.05$, **: $p<0.01$, ***: $p<0.001$, ****: $p<0.0001$.

(E) Non-metric multidimensional scaling (NMDS) analysis of KO profiles was performed to visualize similarities.

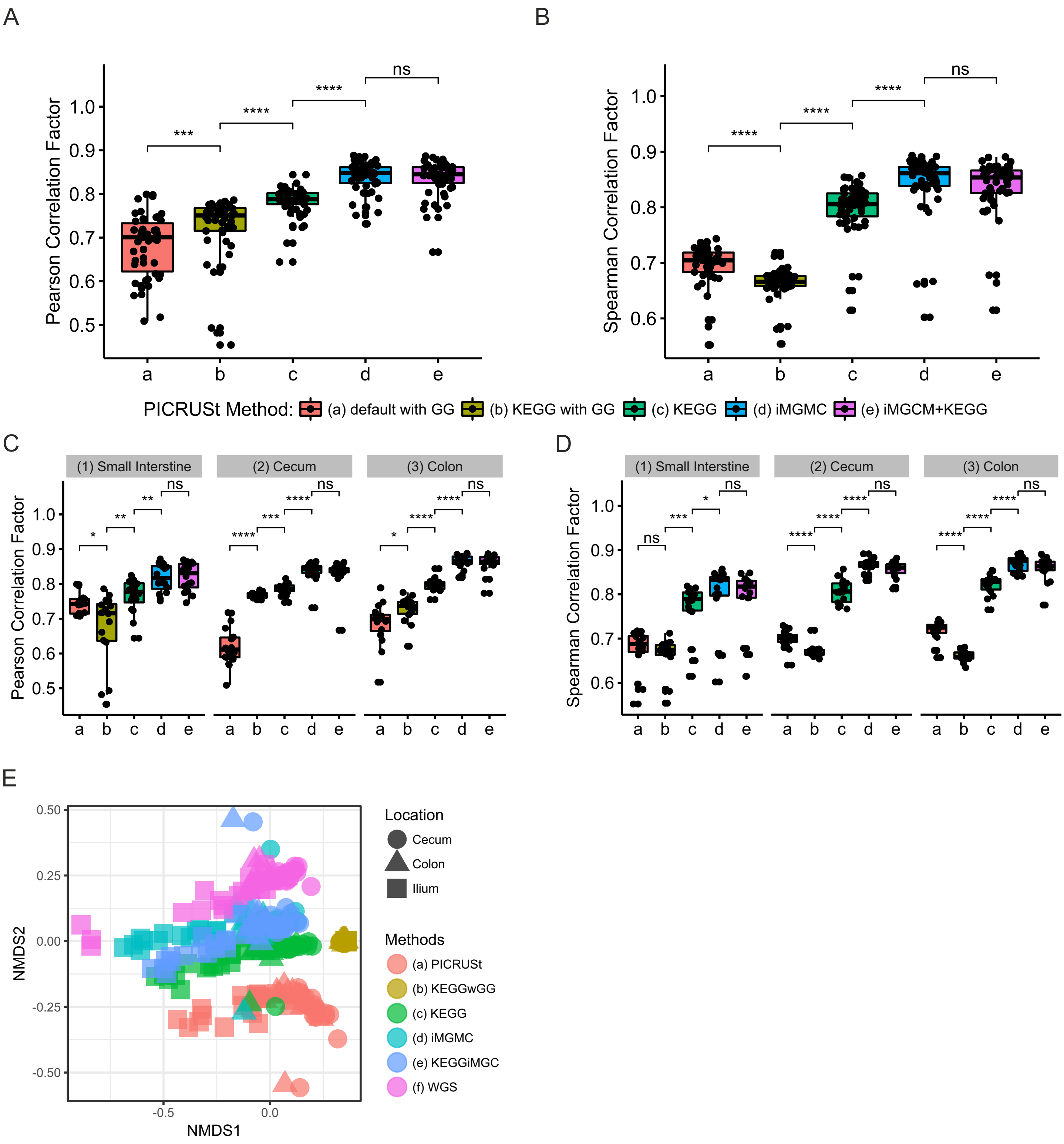
A
B
Pearson Correlation Factor
Spearman Correlation Factor
***
****
ns
a
b
c
d
e
PICRUSt Method: (a) default with GG (b) KEGG with GG (c) KEGG (d) iMGMC (e) iMGCM+KEGG
C
D
(1) Small Interstine
(2) Cecum
(3) Colon
E
Location
Cecum
Colon
Ilium
Methods
(a) PICRUSt
(b) KEGGwGG
(c) KEGG
(d) iMGMC
(e) KEGGiMGC
(f) WGS
NMDS1
NMDS2

Figure-S6

**Figure S7: Evaluation of mouse microbiota optimized PICRUSt model built from 488 iMGMC-MAGs with unique 16S rRNA sequences from external fecal samples**

Correlation between KO profiles of metagenomes (n=15, fecal samples not used to create gene catalog) determined by WGS and different PICRUST workflows using Pearson (A) and Spearman correlation (B), respectively. Correlations factor were determinate for different mouse vendors separately (C and D). Two-tailed paired t-test was performed to analyze the differences, *: p<0.05, **: p<0.01, ***: p<0.001, ****: p<0.0001.

(E) Non-metric multidimensional scaling (NMDS) analysis of KO profiles was performed to visualize similarities.

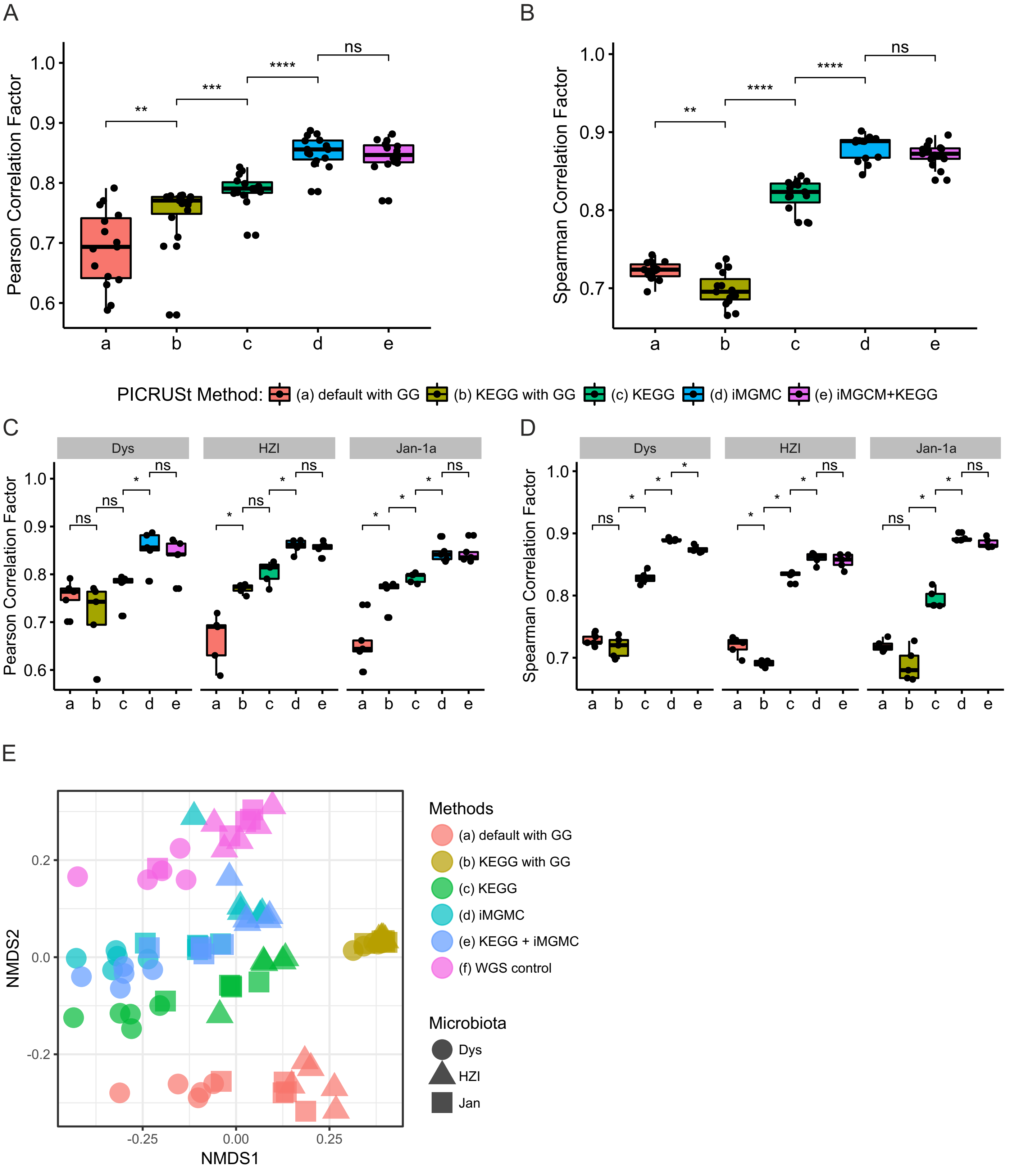
A
B
Pearson Correlation Factor
Spearman Correlation Factor
PICRUSt Method: (a) default with GG (b) KEGG with GG (c) KEGG (d) iMGMC (e) iMGCM+KEGG
C
D
Dys
HZI
Jan-1a
E
NMDS1
NMDS2
Methods
(a) default with GG
(b) KEGG with GG
(c) KEGG
(d) iMGMC
(e) KEGG + iMGMC
(f) WGS control
Microbiota
Dys
HZI
Jan

Figure-S7

**Figure S08: Evaluation of iMGMC using external mouse gut microbiota datasets.** Comparison of mapping rates (A) and taxonomic assignment (B) of previously published mouse gut metagenome datasets based on the original mouse gut catalog (MGCv1, red) and iMGMC (green). Relative improvements of mapping rates are indicated in (A). Two-tailed paired t-test was performed to analyze the differences in (A), ***: $p<0.001$, **: $p<0.01$. Suez et al, 2014, fecal samples (n=40)[3]; Everard et al, 2014, cecal samples (n=34)[4]; Levy et al, 2015, fecal samples (n=10)[5].

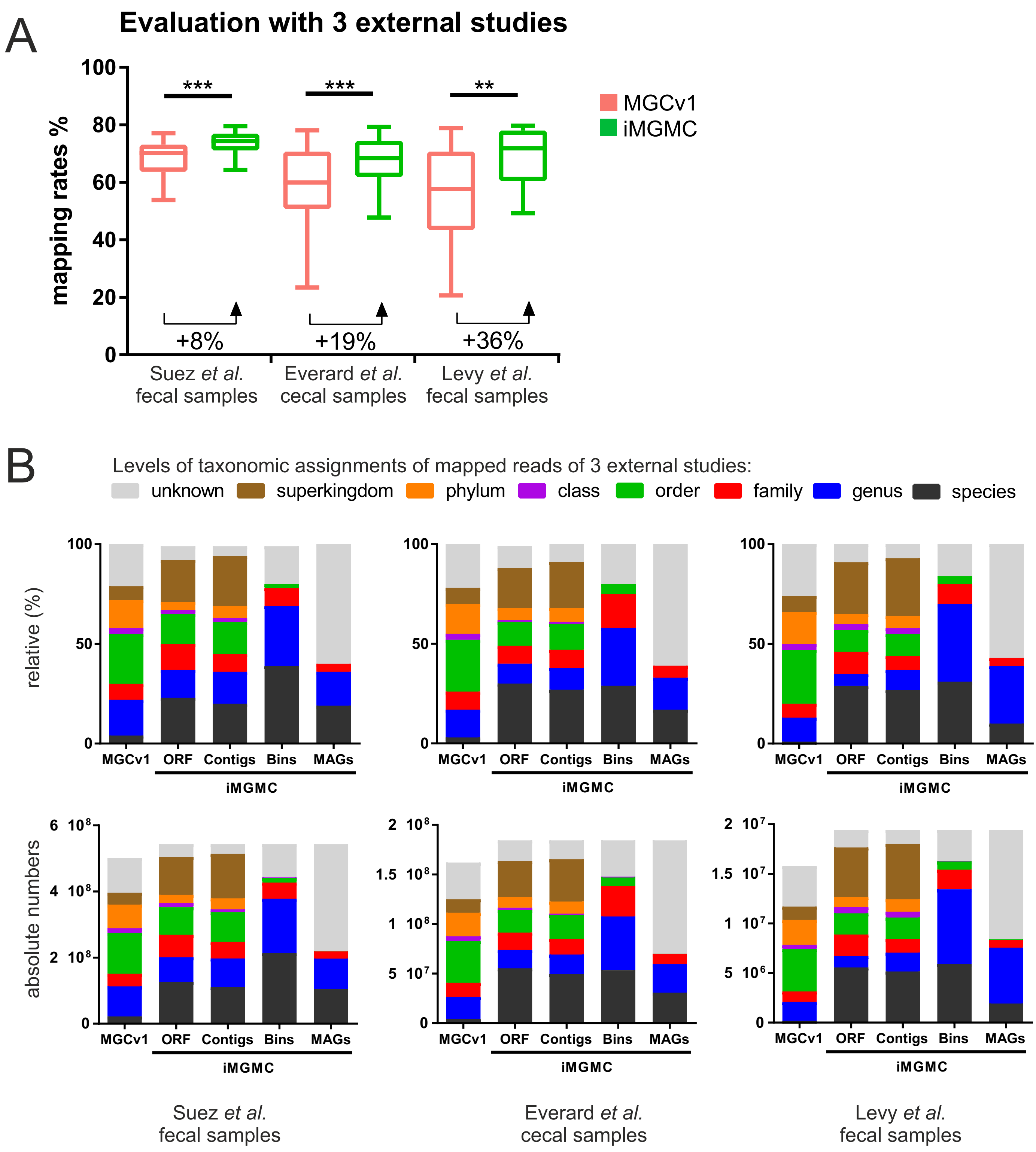
A
Evaluation with 3 external studies
mapping rates %
100
80
60
40
20
0
***
***
**
MGCv1
iMGMC
+8%
+19%
+36%
Suez et al. fecal samples
Everard et al. cecal samples
Levy et al. fecal samples
B
Levels of taxonomic assignments of mapped reads of 3 external studies:
unknown
superkingdom
phylum
class
order
family
genus
species
relative (%)
absolute numbers
MGCv1
ORF
Contigs
Bins
MAGs
iMGMC
Suez et al. fecal samples
Everard et al. cecal samples
Levy et al. fecal samples

Figure S8

**Figure S9: Identification of MAGs networks and their diet induced changes using iMGMC.**

(A) Sub-communities of MAGs were identified by mapping metagenome libraries (N=298, used to generate iMGMC) to MAGs after which the abundance of each MAGs was expressed in TPM and their co-abundances were determined after shrinkage analysis of the correlation factors (see methods for details). Sub-communities with at least 5 members are depicted. Colors of nodes indicate the taxonomic classification of the MAGs and the numbers represent their iMGMC binID (see Table S2). Providers that contain the sub-communities are listed.

(B) Characterization of the influence of diet on sub-communities. Samples from the providers Tac-DK and Jax-US were divided according to the diet and the abundance of individual members of sub-communities 2 and 4 was determined (chow: red; high-fat diet (HF): blue).

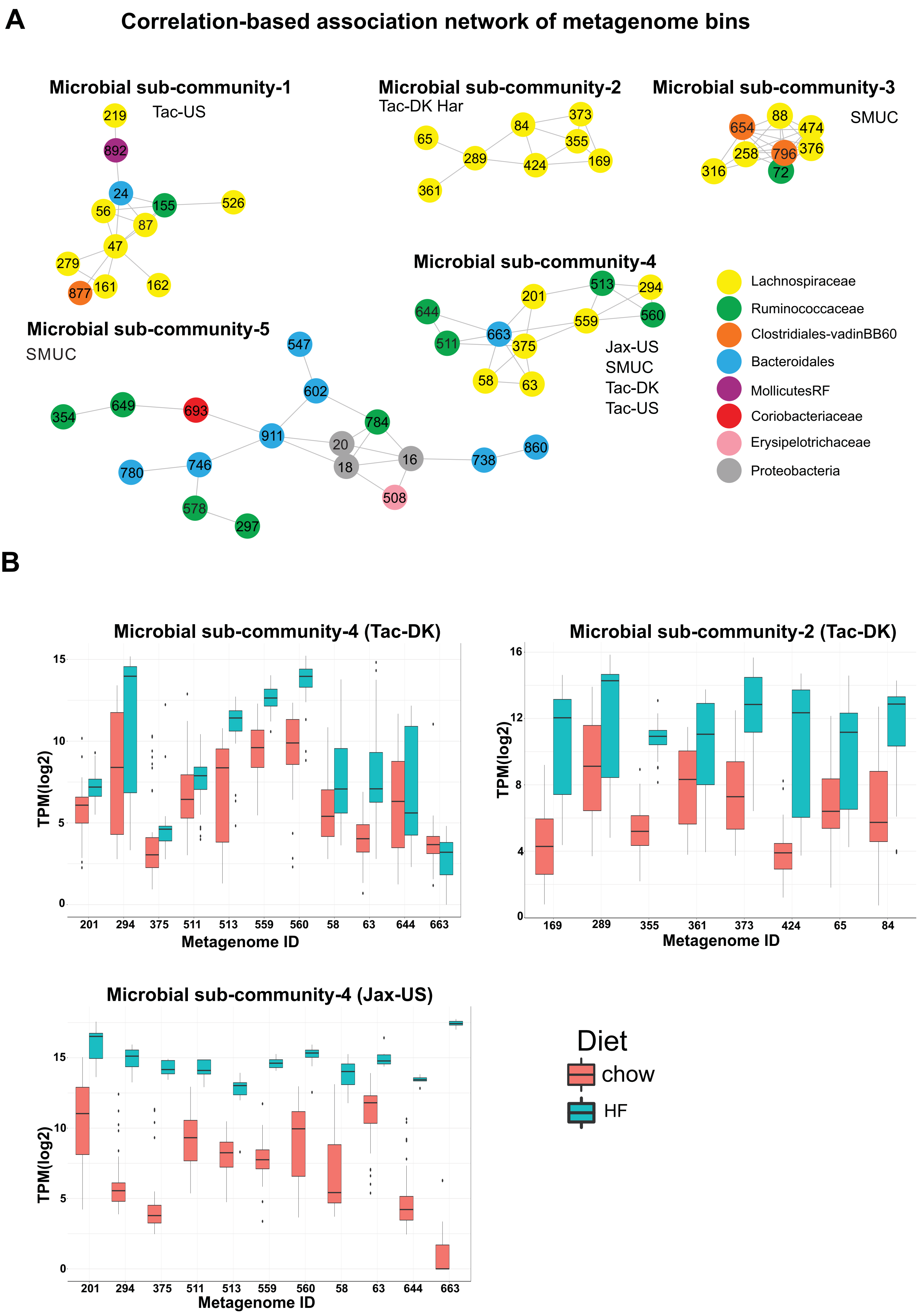
A
Correlation-based association network of metagenome bins
Microbial sub-community-1
Tac-US
Microbial sub-community-2
Tac-DK Har
Microbial sub-community-3
SMUC
Microbial sub-community-4
Jax-US
SMUC
Tac-DK
Tac-US
Microbial sub-community-5
SMUC
Lachnospiraceae
Ruminococcaceae
Clostridiales-vadinBB60
Bacteroidales
MollicutesRF
Coriobacteriaceae
Erysipelotrichaceae
Proteobacteria
B
Microbial sub-community-4 (Tac-DK)
Microbial sub-community-2 (Tac-DK)
Microbial sub-community-4 (Jax-US)
TPM(log2)
Metagenome ID
Diet
chow
HF

Figure-S9

### Material and methods

#### Data availability

All relevant data sets and pipeline scripts are accessible via github.com/tillrobin/iMGMC. Raw reads are available on request.

#### Sample collection and DNA extraction

For the *de novo* generation of metagenomic sequencing data, luminal fecal content was collected from different gastrointestinal GI sites (Ileum: SI, Cecum: Cec and Colon: Col) of mice obtained from different vendors and stored at -20 ˚C until processing. DNA was isolated using an established protocol[6]. Briefly, each sample was treated with 500 µl of extraction buffer (200 mM Tris, 20 mM EDTA, 200 mM NaCl, pH 8.0), 200 µl of 20% SDS, 500 µl of phenol:chloroform:isoamyl alcohol (24:24:1) and 100 µl of zirconia/silica beads (0.1 mm diameter). Samples were homogenized with a bead beater (BioSpec) for 2 min. DNA was precipitated with absolute isopropanol and finally washed with 70 % ethanol. DNA extracts were resuspended in TE Buffer with 100 µg/ml RNase I and finally column purified to remove traces of PCR inhibitors.

#### Metagenomic sequencing

Total DNA was quantified and diluted to 25 ng/µl. 60 µl of total DNA were used for shearing by sonication (Covaris). Fragmentation was performed as follow: Processing time = 150 s, Fragment size = 200 bp, Intensity= 5, duty cycle= 10. Illumina library preparation was performed using the NEBNext Ultra DNA library prep kit (New England Biolabs). The library preparation was performed according to the manufacturer's instructions. We use as input a total of 500 ng of DNA, the size selection was performed using AMPure XP beads (First bead selection = 55 µl, and second = 25 µl). Adaptor enrichment was performed using seven cycles of PCR using the NEBNext Multiplex oligos for Illumina (Set 1 and Set 2)(New England Biolabs) and then subjected to Illumina HiSeq2000 PE100 sequencing. Source and sequencing depth for each sample is listed in table S1.

#### 16S rRNA gene amplification, sequencing and data analysis

Amplification of the V4 region (F515/R806) of the 16S rRNA gene was performed according to previously described protocols [7,8]. Briefly, for DNA-based amplicon sequencing 25 ng of DNA were used per PCR reaction (30 µl). The PCR conditions consisted of initial denaturation for 30s at 98°C, followed by 25 cycles (10s at 98°C, 20s at 55°C, and 20s at 72°C. Each sample was amplified in triplicates and subsequently pooled. After normalization PCR amplicons were sequenced on an Illumina MiSeq platform (PE250). Obtained reads were assembled, quality controlled and clustered using the QIIME v1.8.0 (Quantitative Insights Into Microbial Ecology) analysis pipeline [9]. In short, quality filtering was set to -q 30, minimum read length 200 bp and minimum number of sequences per sample = 1000. The OTU clusters and representative sequences were determined using open-reference OTU picking [10] using UCLUST [11] at 97% identity, followed by taxonomy assignment using the RDP Classifier [12] with a bootstrap confidence cutoff of 80%. The OTU absolute abundance table and mapping file were used for statistical analyses and data visualization in the R statistical programming environment (R Core Team (2016). R: A language and environment for statistical computing. R Foundation for Statistical Computing, Vienna, Austria. URL https://www.R-project.org/.)

**Construction of the iMGMC**

**i) Assembly and prediction of ORFs**

Demultiplexed libraries were filtered to remove host reads using bbmap (parameters see code) using the Ensembl masked mouse genome GRCm38.75 and phiX. All mouse filtered metagenomic libraries were used in single “all in one” assembly approach using Megahit[13] with specific parameters (-kmin 5 -k 27,37,47,57,67,77,87,97) using a SGI-UV2000 cluster with 256 cores and 2 TB shared memory. Resulting contigs were filtered to minimum 1000bp lengths and renamed with numbers from largest to smallest contig. For protein prediction we used Metagenemark [14] (parameters see code). ORFs were filtered to remove ORF shorter than 100bp after which they were reordered and renamed according to their length (Fig 1A, 1D).

**ii) Binning and evaluation of binning using CheckM**

All libraries were mapped with BWA [15] (default parameters) to the contigs. The mapping results were transformed and indexed to bam-format using sambawa. Metagenome binning was performed with MetaBAT [16](version 0.32) using the following parameters -verysensitive -pB 20 -B 100 -minclustersize 200000. The resulting clusters were evaluated with CheckM [17]. To assign a bin to a high quality MAGs (metagenome species) we used a threshold of marker gene completeness - contamination >= 80%. All other bins of contigs with at least 200kbp length were defined as CAGs (Co-abundance groups). We used the marker gene alignment from CheckM derived from all 660 MAGs and 64 selected genomes from NCBI RefSeq to construct a phylogenetic tree using a nearest neighbor joining approach with 1000 bootstraps in MEGA7[18]. Tree was plotted using GraPhlAn[19].

**iii) Taxonomic classification**

Taxonomic classification for all gene entries in the catalog was performed based on different levels, i.e. ORF, contig and bin/CAG/MAGs. For ORFs, assignments were performed using

CAT and DIAMOND [20] against the NCBI NR protein database. For all contigs and bins the classification was performed by taxator, ppsp, GTDB-Tk and contig annotation tool CAT using default parameters. We use conducted contigs for the bin assignments.
Furthermore we used clustering of the MAGs-Tree and 16S linking with Silva taxonomy to make a manual annotations of the MAGs.

**iv) Functional annotation of gene catalog proteins**

All proteins were annotated using blastp[21] against the KEGG gene database (01/2018) [22] following the best hit approach (evalue 0.001). KEGG Orthology annotation was used to reconstruct KEGG module completeness. For the annotation for the MAGs we using linked data form catalogue. Libraries were mapped to gene catalog, all KO genes which have one mapped read were counted as present.We used for statistical analyses and data visualization in the R statistical programming environment.

**v) Full length 16S rRNA sequence reconstruction, annotation and phylogeny**

We were used RAMBL[23] to reconstruct full length 16S rRNA gene sequences from all libraries in one batch. Resulting sequences (n = 1323) were classified with SINA[24] using the SILVA NRref database (version 123). Further on sequences were processed by Blast search against NCBI16Sref database to find reference 16S sequences and genomes. A phylogenetic tree built by nearest neighbour joining method (maximum likelihood) with 1000 bootstraps using MEGA7[18].

**vi) MAG to 16S rRNA gene connections via multi-scale linkage**

The full linking pipeline is based on three different approaches: First, we searched for integrated 16S rRNA sequences in the assembled contings of all clustered bins (CAG/MAGs). For this we mapped with BlastN all contigs to all reconstructed 16S rRNA gene sequences. We filter out alignments of less than 100bp and lower than 95% of identity, resulting in a matrix of blast scores of each bin to each 16S rRNA gene sequence.
Second, we used a scaffolding approach using paired-end read information. The reads from all libraries were partitioned into new libraries by mapping against all bins (n=1462) with bbsplit (Ref bbmap). Then, the new libraries were mapped against all 1,323 reconstructed 16S sequences. Unambiguous and ambiguous mapped reads were counted separately into two matrices of all bins x 16S rRNA gene sequences.
The third method uses the abundance profiles across all samples to correlate 16S rRNA gene sequences and bins. To determine abundance profiles over all 298 samples, we mapped all libraries individually to all CAG/MAGs (and unbinned contigs) and to all reconstructed full-length 16S rRNA gene sequences. MAGs read counts were transformed to TPM (transcripts

per million) and stored in an abundance matrix, as well the unambiguous 16S rRNA gene sequence counts. Pearson and Spearman correlation were calculated for both abundance matrices to get scoring for all bins to all 16S rRNA gene sequences.

Finally, data of all three approaches were weighed in an integration scoring to aim in associating the meta-genomic bins of the mouse catalogue to the corresponding 16S annotations, in different steps:

*(I) Indirect association:* We used the normalized abundance values of the metagenome bins and 16S rRNA gene sequence data to obtain their corresponding correlation (both Pearson and Spearman). We estimated consensus interdependence score from both the correlation methods between any 16S rRNA gene and metagenome bin pair by integrating correlation values between the metagenome bins and 16S rRNA genes by taking the geometric mean of both the correlation values between each metagenome bin and 16S rRNA gene and assigning a negative sign if either of these correlation values was negative.

$$V(x,y) = value[I(x,y)] = \sqrt{abs\big(P(x,y)\big) * abs(S(x,y))}$$

$$Sg(x,y) = sign\big[\big(I(x,y)\big)\big] = \begin{cases} -, & any\ [P(x,y), S(x,y)] < 0 \\ +, & else \end{cases}$$

$$I(x,y) = V(x,y) * Sg(x,y)$$

Where

$P(x,y)$= Pearson correlation between a 16S rRNA gene ‘x’ and metagenome bin ‘y’

$S(x,y)$= Spearman correlation between a 16S rRNA gene ‘x’ and metagenome bin ‘y’

$I(x,y)$= Integrated correlation between a 16S rRNA gene ‘x’ and metagenome bin ‘y’

For the highly correlated metagenome-16S rRNA gene pairs, the Pearson and Spearman correlation values have very small difference. The normalization described above widened the distance between the true positives and false positives.

*(II) Direct association:*

1. Mapping bins to 16S rRNA gene sequences [$M(x,y)$]: These quantify the fraction of reads in a bin ‘y’ containing matching reads from 16S rRNA gene ‘x’ by mapping the reads in bin ‘y’ to the 16S rRNA gene ‘x’. We normalized the number of uniquely mapped reads in bin ‘y’ to a 16S rRNA gene ‘x’ $m(x,y)$ by the total number of 16S reads mapped to the bin ‘y’ [$\sum_{i=1}^{n} m(x,i)$].

   $$\mathrm{M}(x,y) = \frac{m(x,y)}{\sum_{i=1}^{n} m(x,i)}$$

   Where n is the number of bins

2. BLAST bins to 16S rRNA gene sequences [$B(x,y)$]: These quantify the fraction of reads in a bin ‘y’ containing reads in rRNA gene ‘x’ aligning of the reads of rRNA

gene ‘x’ to the bin ‘y’ using BLAST. We normalized the number of uniquely mapped reads in bin ‘y’ to a rRNA gene ‘x’ [$b(x, y)$] by the maximum of reads from rRNA genes mapped to the bin ‘y’ [$\max_{0<i\le n} b(x, i)$].

$$\mathrm{B}(x, y) = \frac{b(x, y)}{\max\limits_{0<i\le n} b(x, i)}$$

Where n is the number of metagenome bins.

*(III) Integrating the direct and indirect associations between bin and 16S rRNA gene sequences:*

The direct associations are sparse, i.e. there are very few 16S rRNA gene sequences reads present in each bin, while the indirect associations are not sparse. Hence, we integrated the scores in a way that does not allow the indirect associations to dominate over the direct associations. For this, we integrated the three scores [I(x,y), $M(x, y), B[(x, y)]$, as done in the STRING database. The only difference between the STRING database and the scoring scheme employed here is that for combining scores we took a geometric mean of the dissimilarity scores while combining them instead of simply multiplying the different scores (as done in STRING database).

$$F = 1 - \sqrt[3]{(1 - I) * (1 - B) * (1 - M)}$$

Where F is the combined score for the bins – 16S rRNA gene sequences relationship. We observed that the integrated correlation scores [$I(x, y)$] representing the indirect association tended to dominate over the direct association scores in several instances. Hence, we regularized the indirect association score by multiplying the Pearson and Spearman correlation values, instead of calculating their geometric mean:

$$V_{reg}(x, y) = value\left[I_{reg}(x, y)\right] = abs\left(P(x, y)\right) * abs(S(x, y))$$

$$I_{reg}(x, y) = V_{reg}(x, y) * \ Sg(x, y)$$

$$F_{reg} = 1 - \sqrt[3]{\left(1 - I_{reg}\right) * (1 - B) * (1 - M)}$$

Where

$V_{reg}$: Regularized integrated correlation value.

$I_{reg}$ : Regularized integrated correlation score.

$F_{reg}$ : Regularized combined score for the metagenome-rRNA genes relationship.

The negative values are turned zeros. The closer the $F_{reg}$ value to 1, the higher the confidence of the bin – 16S rRNA gene sequences relationship. However, the 16S rRNA gene sequence ‘x’ might have the highest confidence score to the metagenome bin ‘y’, but the metagenome bin ‘y’ need not have the highest confidence score to rRNA gene ‘x’. To address this issue, we enriched these relationships by normalizing these scores by the highest confidence scores of the corresponding metagenome bin ’y’ and rRNA gene ‘x’.

*(IV) Enriching metagenome bin to rRNA gene relationship:*

We estimated the probability of a metagenome bin ‘y’ to rRNA gene ‘x’ relationship:

$$\Pr(x,y) = \frac{F_{reg}(x,y)}{\max\limits_{0<i\leq m} F_{reg}(i,y)} * \frac{F_{reg}(x,y)}{\max\limits_{0<j\leq n} F_{reg}(x,j)}$$

Where n is the number of bins and m is the number of 16S rRNA genes.

The obtained normalized confidence score or the estimated probability is the statistical likelihood of the confidence scores, adjusted for the background distribution of the confidence scores for all possible 16S-metagenome pair relationships.

**Evaluation of iMGMC:**

**i) Assessing binning efficiency using know NCBI reference genomes recovered as MAGs**

In order to evaluate the binning of contigs of higher order, those reference genomes being contained within the assembly were identified. Therefore, synthetic reads (100 bp) from all 9,748 bacterial genomes available in the NCBI Assembly database (Version January 2017) were generated with BBMap and mapped against all contigs. Genomes that were contained to at least 50% within the contigs were selected for evaluation (n = 57). Specifically, the binning efficiency for each reference genome was evaluated by quantifying the distribution of the synthetic reads over the bins and unbinned contigs. The analysis contained both the total proportion of reads mapped to contigs (= total recovered genome fraction) as well as the fraction of reads contained within contigs and mapping to one or more bins (= binned genome fraction).

ii) **Assessing the bins to 16S rRNA gene linking approach with reference genomes**

In order to evaluate the linking approach, those NCBI reference genomes which constitute part of MAGs were identified. Therefore, synthetic reads (100 bp) from all 9,748 bacterial genomes available in the NCBI Assembly database (Version January 2017) were generated with BBMap and mapped against all MAGs. Those mapping to at least 50% to a single MAGs, were used for evaluating the MAG / 16S rRNA gene links. First, the 16S rRNA gene sequences of the NCBI genomes were matched to the best-reconstructed 16S rRNA gene sequences via BlastN and the identify was calculated. Then, these reference sequences were compared to the predicted 16S rRNA gene sequence from the linking approach and the taxonomic agreement between these sequences was scored. Optimally, an identical match of reference

16S rRNA gene sequence to the linked 16S rRNA gene sequence would obtained by the scoring scheme.

#### PICRUSt

To test if the MAGs linked with reconstructed 16S rRNA sequences represent a large part of the mouse gut catalogue we created an extended genomic reference PICRUSt prediction model: 484 MAGs with unique linked 16S sequences were used according the PICRUSt "Genome Prediction Tutorial": 1) Determination of 16S copy numbers was performed by rrnDB Estimate (version 5.2.), 2) KEGG Orthology (KO) profiles of the MAGs were extracted from iMGMC. 3) A tree [25,26] of RAMBL reconstructed 16S sequences was used to build the models. Furthermore, to verify the prediction power of the model we added the KO profiles of 3772 KEGG genomes to iMGMC model. Moreover, use the sequences of the GreenGenes database (Version 13.5 OTU-RepSet 97) together with the iMGMC reconstructed 16S sequences and the 16S of the KEGG genomes to build an integrated prediction model.
To make our PICRUSt models accessible for de-novo clustered OTUs, we modified our pipeline in a way that for each dataset, a new PICRUSt model is created from stretch: Sequences of the OTUs where include by pynast[27] into a precalculated alignment of the reconstructed 16S rRNA genes. A tree generated by FastTree[26] is be used for the PICRUSt genome predictions to create a pre-calculated PICRUSt models that include the denovo OTUs for the mapping into the following standard workflow.

#### Global distribution of iMGMC 16S rRNA gene sequences in NCBI-SRA (IMNGS analysis)

We checked all 1,323 reconstructed full-length 16S rRNA gene sequences using the IMNGS pipeline[28] for their prevalence and relative abundance in 168,573 SRA samples (build 1711). To evaluate the specific environment of an OTU, we looked also at its relative abundance in related mouse gut environments such as the skin (mouse, human) and different gut sites (human, rat, bovine, chicken, fish, insect, pig, termite) where at least 100 samples were available. We filter for 16S rRNA less than 0.1% abundance in one of the selected environments. Furthermore, we checked for specific preference of an OTU in mouse gut, human gut and rat gut, by using a relative abundance normalized over the environments of at least 50%. Other OTUs where moreover checked to be dominant in combination with the mouse gut site together with mouse skin, human gut or rat gut.

#### Identification of sub-communities in the intestinal bacterial community

We obtained the co-abundances between all metagenome bin-pairs using a shrinkage approach to the correlation estimation, as described in [29] (and available of cor.shrink function in corpcor library in R). We removed the co-abundance values less than 0.5 and inferred the sub-communities using a modularity optimization algorithm described in [30] (and available as cluster_louvain function in igraph library in R). Amongst all the sub-communities obtained, we were interested in those sub-communities of more than 5 members.

Association between sub-communities and diet across multiple vendor mouse gut-microbial communities:

We performed multi-factor ANOVA address relations of difference in dietary supplements and the difference in mouse strains over the abundance of the metagenome bins. For this, we modelled the metagenome bins abundance data against the interaction effect of the metagenome bin and dietary supplements, the dietary supplements and the mouse strains of samples from every vendor separately. Any sub-community which showed a significant response to difference in diet, metagenome bin and the interaction effect of diet and the metagenome bin (p-value less than 1%), but not a significant response to difference in mouse strains (p-value more than 10%) was considered to have an association between diet interventions and the microbial sub-community.

**References:**


1. Mikheenko, A., Saveliev, V. & Gurevich, A. MetaQUAST: evaluation of metagenome assemblies. *Bioinformatics* **32,** 1088–1090 (2016).
2. Alneberg, J. *et al.* Binning metagenomic contigs by coverage and composition. *Nat. Methods* **11,** 1144–1146 (2014).
3. Suez, J. *et al.* Artificial sweeteners induce glucose intolerance by altering the gut microbiota. *Nature* **514,** 181–186 (2014).
4. Everard, A. *et al.* Microbiome of prebiotic-treated mice reveals novel targets involved in host response during obesity. *ISME J.* **8,** 2116–2130 (2014).
5. Levy, M. *et al.* Microbiota-Modulated Metabolites Shape the Intestinal Microenvironment by Regulating NLRP6 Inflammasome Signaling. *Cell* **163,** 1428–1443 (2015).
6. Turnbaugh, P. J. *et al.* A core gut microbiome in obese and lean twins. *Nature* **457,** 480–484 (2009).
7. Caporaso, J. G. *et al.* Global patterns of 16S rRNA diversity at a depth of millions of sequences per sample. *Proc. Natl. Acad. Sci.* **108,** 4516–4522 (2011).
8. Thiemann, S. *et al.* Enhancement of IFNγ Production by Distinct Commensals Ameliorates Salmonella -Induced Disease. *Cell Host Microbe* **21,** 682–694.e5 (2017).
9. Caporaso, J. G. *et al.* QIIME allows analysis of high-throughput community sequencing data. *Nat. Methods* **7,** 335–336 (2010).
10. Nelson, M. C., Morrison, H. G., Benjamino, J., Grim, S. L. & Graf, J. Analysis, optimization and verification of Illumina-generated 16S rRNA gene amplicon surveys. *PLoS One* **9,** e94249 (2014).
11. Edgar, R. C. Search and clustering orders of magnitude faster than BLAST. *Bioinformatics* **26,** 2460–2461 (2010).
12. Wang, Q., Garrity, G. M., Tiedje, J. M. & Cole, J. R. Naive Bayesian Classifier for Rapid Assignment of rRNA Sequences into the New Bacterial Taxonomy. *Appl. Environ. Microbiol.* **73,** 5261–5267 (2007).
13. Li, D. *et al.* MEGAHIT v1.0: A fast and scalable metagenome assembler driven by advanced methodologies and community practices. *Methods* **102,** 3–11 (2016).
14. Zhu, W., Lomsadze, A. & Borodovsky, M. Ab initio gene identification in metagenomic sequences. *Nucleic Acids Res.* **38,** e132–e132 (2010).
15. Li, H. & Durbin, R. Fast and accurate short read alignment with Burrows-Wheeler transform. *Bioinformatics* **25,** 1754–1760 (2009).
16. Kang, D. D., Froula, J., Egan, R. & Wang, Z. MetaBAT, an efficient tool for accurately reconstructing single genomes from complex microbial communities. *PeerJ* **3,** e1165 (2015).

17. Parks, D. H., Imelfort, M., Skennerton, C. T., Hugenholtz, P. & Tyson, G. W. CheckM: assessing the quality of microbial genomes recovered from isolates, single cells, and metagenomes. *Genome Res.* **25,** 1043–1055 (2015).
18. Kumar, S., Stecher, G. & Tamura, K. MEGA7: Molecular Evolutionary Genetics Analysis Version 7.0 for Bigger Datasets. *Mol. Biol. Evol.* **33,** 1870–1874 (2016).
19. Asnicar, F., Weingart, G., Tickle, T. L., Huttenhower, C. & Segata, N. Compact graphical representation of phylogenetic data and metadata with GraPhlAn. *PeerJ* **3,** e1029 (2015).
20. Buchfink, B., Xie, C. & Huson, D. H. Fast and sensitive protein alignment using DIAMOND. *Nat. Methods* **12,** 59–60 (2015).
21. Altschul, S. F. *et al.* Gapped BLAST and PSI-BLAST: a new generation of protein database search programs. *Nucleic Acids Res.* **25,** 3389–402 (1997).
22. Kanehisa, M., Furumichi, M., Tanabe, M., Sato, Y. & Morishima, K. KEGG: new perspectives on genomes, pathways, diseases and drugs. *Nucleic Acids Res.* **45,** D353–D361 (2017).
23. Zeng, F., Wang, Z., Wang, Y., Zhou, J. & Chen, T. Large-scale 16S gene assembly using metagenomics shotgun sequences. *Bioinformatics* **33,** 1447–1456 (2017).
24. Pruesse, E., Peplies, J. & Glöckner, F. O. SINA: Accurate high-throughput multiple sequence alignment of ribosomal RNA genes. *Bioinformatics* **28,** 1823–1829 (2012).
25. Edgar, R. C. MUSCLE: multiple sequence alignment with high accuracy and high throughput. *Nucleic Acids Res.* **32,** 1792–1797 (2004).
26. Price, M. N., Dehal, P. S. & Arkin, A. P. FastTree 2 – Approximately Maximum-Likelihood Trees for Large Alignments. *PLoS One* **5,** e9490 (2010).
27. Caporaso, J. G. *et al.* PyNAST: a flexible tool for aligning sequences to a template alignment. *Bioinformatics* **26,** 266–267 (2010).
28. Lagkouvardos, I. *et al.* IMNGS: A comprehensive open resource of processed 16S rRNA microbial profiles for ecology and diversity studies. *Sci. Rep.* **6,** 33721 (2016).
29. Schäfer, J. & Strimmer, K. A Shrinkage Approach to Large-Scale Covariance Matrix Estimation and Implications for Functional Genomics. *Stat. Appl. Genet. Mol. Biol.* **4,** Article32 (2005).
30. Blondel, V. D., Guillaume, J.-L., Lambiotte, R. & Lefebvre, E. Fast unfolding of communities in large networks. *J. Stat. Mech. Theory Exp.* **2008,** P10008 (2008).
